# Sustainability or Otherwise: The Lotka-Volterra Model in Population Ecology and Socio-Economic Systems

**DOI:** 10.1101/2025.05.01.651785

**Authors:** N. Karjanto, B. Sriraman

## Abstract

The Lotka-Volterra equations, a cornerstone of predator-prey modeling, offer a versatile framework for understanding population dynamics in the context of sustainability. This chapter explores how these equations, through the analysis of dynamic equilibrium and stability, illuminate principles of both sustainable ecological interactions and the mechanisms leading to detrimental long-term outcomes in various systems (negative sustainability). We first analyze predator-prey interactions to gain insights into population management, resource conservation, and the delicate balance of ecosystems. Furthermore, we extend the Lotka-Volterra model to discuss its broader implications for complex systems dynamics in diverse fields, such as in modeling economic competition and arms races, where the framework helps identify pathways that lead to unsustainable resource consumption, increased risk, or collapse.

## 1 Introduction

Antelopes and lions, foxes and rabbits, lynx and hares, owls and mice, ladybirds and aphids, dragonflies and mosquitoes–these are just some of the predator-prey pairs commonly found in nature. Each of these relationships is more than a simple predator-prey interaction. The populations of each species are intricately linked, demonstrating the diversity and complexity of ecological interactions.

While this list includes pairs of animals interacting in natural cycles, it often overlooks the most formidable predator of all–humans. Our relentless pursuit for food and resources places us at the top of the predatory chain, with practices like fishing and hunting exerting unparalleled pressure on ecosystems and threatening the balance that sustains them. This raises not only ethical and moral questions about our responsibilities but also crucial sustainability concerns: Can we fish sustainably without depleting marine ecosystems? How might understanding predator-prey dynamics contribute to sustainable resource management? As human activities significantly alter natural ecosystems, understanding these interactions is critical for promoting sustainability both in the wild and in human-modified systems such as agriculture and fisheries.

To answer these sustainability questions, we turn to mathematical modeling, which provides structured insights into predator-prey dynamics. While mathematics excels at revealing systematic patterns, it also has limitations in capturing the full complexity of ecological systems.

In ecology, the interaction between predator and prey falls under the concept of *biocoenosis*. Also often referred as *life assemblage* or *biotic community*, it refers to an association of different species cohabiting in the same area, which we refer to as a *biotope*, at the same period (von Bertalanffy 1968). In general, many species interact with each other to form structurally complex web of communities, primarily for the purpose of nutrition and survival. A *trophic web*, or *food web*, represents this complex network of energy flow between organisms in an ecosystem, highlighting the interdependence among species (Murray 2007).

Using a stock-flow diagram, we can visualize the dynamics of predator-prey interactions, as illustrated in Figure 1 (Roe et al. 2018). In this diagram, green rectangles represent stocks, denoting the predator and prey populations, while blue ovals indicate the inflows (birth rates) and outflows (death rates) affecting each population.

**Figure 1.**
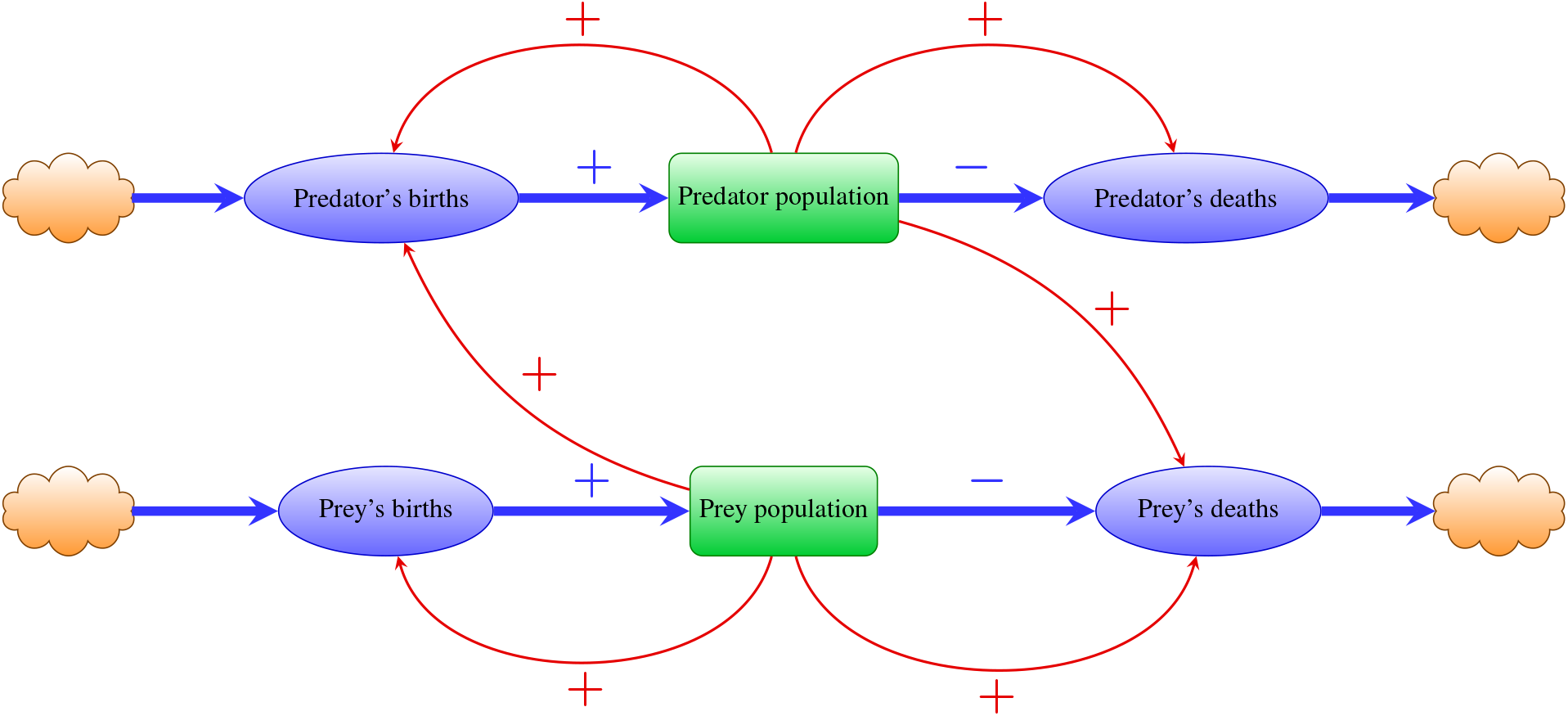
An interaction between predator and prey populations depicted as a stock-flow diagram.

The straight blue arrows indicate inflow and outflow rates, with positive arrows for births and negative arrows for deaths. The clouds at both ends of the flow represent sources and sinks, assuming infinite capacity, which ensures no external constraints affect the flow.

The inflow rate of natality and the outflow rate of mortality for each species are affected by amplifying and stabilizing feedback loops, respectively. These are indicated by the curved red arrows with positive sign that emerge from the stock of predator and prey populations to the inflow of birth rates and the outflow of death rates. Additionally, there are positive control effects of the prey population on the predator’s natality rate and the predator’s population on the prey’s mortality rate. These feedback loops play a crucial role in sustainability studies, as they enable us to predict how changes in one population affect the entire ecosystem.

A mathematical model for the predator and prey relationship has been long known, with the Lotka-Volterra equations being the notable one. This model describes how predator and prey populations interact over time. The equations capture the growth of the prey population and the dependency of the predator population on prey availability, leading to cyclical population dynamics.

This model was proposed by a Polish-American mathematician Alfred James Lotka in the 1910s and 1920s (Lotka 1920, Lotka 1925). In 1926, an Italian mathematician Vito Volterra published the same set of equations and credited Lotka’s earlier work (Volterra 1926a, Volterra 1926b, Volterra 1928, Volterra 1931). The model that involves two species became known as the “classical Lotka-Volterra model” since then. This model can help us understand the dynamics of predator-prey populations and how they oscillate and evolve over time.

However, real ecosystems often involve complex food webs with multiple species interactions. This model, while foundational, may oversimplify these dynamics, limiting its applicability to more intricate ecosystems (Brauer 2012, Hannon and Ruth 2014). Nevertheless, the enduring theoretical importance of the Lotka-Volterra framework, including discussions and generalizations of principles such as the Volterra principle, continues to be explored in modern scholarship (Räz 2017).

In this chapter, we delve into how the Lotka-Volterra predator-prey model serves as a powerful tool for understanding sustainability, both in promoting sustainable ecological practices and in identifying dynamics that lead to detrimental, or “negative,” long-term outcomes across various systems. This chapter is organized as follows. Following this introduction, Section 2 delves into the mathematical foundations of the classical Lotka-Volterra predator-prey model, detailing its exact solutions, properties, and the characteristics of its population equilibria.

Section 3 then explores the multifaceted relationship between the model’s dynamics and sustainability. While demonstrating how understanding cyclical population dynamics can inform sustainable resource management and harvesting strategies, it crucially extends the framework to non-ecological competitive scenarios, such as arms races, to illuminate the concept of “negative sustainability”— where inherent system dynamics can lead to unsustainable resource consumption, increased risk, or collapse. Finally, Section 4 provides a comprehensive summary and concluding remarks.

## 2 The Lotka-Volterra Model

The classical Lotka-Volterra equations consist of a system of first-order nonlinear ordinary differential equations (ODEs) that describes the interaction of two species, one being a predator, and the other as prey in a biocoenotic system. Let *N* (*t*) be the prey population and *P* (*t*) be the predator population, and both depend on the time variable *t*. On the one hand, the discrete model treats these populations as a collective of individual units of nonnegative integers and their changes occur at distinct time steps. For example, we can observe the number of individual hares and lynx at the end of each month or year in a particular region where they interact. It follows that in the absence of predators, the prey population can be represented by the following difference equation:

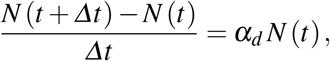

where *Δt* is the time step and *α*_*d*_ is the average per-capita growth rate of the population over discrete intervals. A similar difference equation can also be constructed for the predator population.

On the other hand, when we consider an infinitely small time step, i.e., *Δt* → 0, we can also approach the situation using a continuous model. In this framework, the populations are treated as continuous quantities of densities, allowing for any nonnegative real values and not necessarily integers. Hence, *N* and *P* denote the number of prey and predator animals per unit area on land (or volume in aquatic settings), respectively. Consequently, in the absence of predators, the prey population grows exponentially according to the following differential equation:

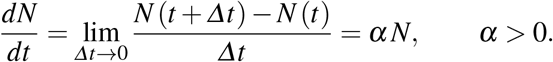

Similarly, we can also transition from the discrete model of a difference equation to the continuous one for the predator population. Without prey, the predator population declines at a rate proportional to itself, and it can be described using the following differential equation:

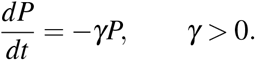

The positive constants *α* and *γ* are called *(instantaneous) natural growth rate* and *(instantaneous) natural decay coefficients*, respectively (Borrelli and Coleman 2004). For the rest of this chapter, we will adopt the continuous model instead of the discrete one.

With both species present, the principal cause of death among the prey is being eaten by a predator. And similarly, the birth and survival rates of the predators depend on their available food supply, i.e., the prey. Assume also that the two species encounter each other at a rate that is proportional to both populations: ∼ *NP*. The term −*β NP, β* > 0, decreases the natural growth rate of the prey, whereas the term +*δ NP, δ* > 0, increases the natural growth rate of the predator. The coefficients *β* and *δ* measure the likelihood that a predator-prey encounter removes prey, and predator efficiency in converting food into fertility, respectively.

This assumption follows the *Population Law of Mass Action*, which states that at time *t*, the rate of change of one population due to interaction with another is proportional to the product of the two populations at that time *t* (Borrelli and Coleman 2004). Furthermore, by incorporating these interaction terms and applying the Balance Law to each population, i.e., the net rate of change of a population = rate in − rate out, we obtain the *predator-prey* (or *Lotka-Volterra*) model, given as follows:

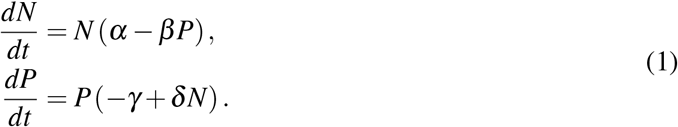

Because the independent variable *t* does not appear explicitly on the right side of each ODE, this Lotka-Volterra model belongs to the class of an *autonomous system* of first-order differential equations (Zill 2023a, Zill 2023b). The model is also nonlinear, thanks to the interactions between the predator and prey.

Although biologically it just makes sense to assume that both *N* and *P* are nonnegative, it can be explained mathematically as well. In the absence of coupling constants *β* and *δ*, the predator and prey will grow and decay exponentially, respectively. However, by coupling both species, the system (1) has the following family of solutions:

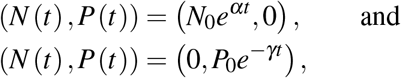

where *N*_0_ > 0 and *P*_0_ > 0 denote the initial prey and predator population density, respectively. For any *t* ≥ 0, both *N* (*t*) and *P* (*t*) > 0, implying that any solution that starts in the first quadrant of the *NP*-plane will remain there for all future time.

The dimensions of the coefficients are given as follows:

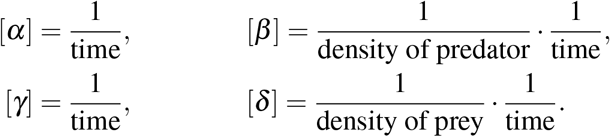

The assumption of the interaction term as the product of the population densities *NP* also assumes a linear increase in consumer’s (in this case, predator’s) intake rate as a function of food (or prey) density. In ecology, this is known as Type I *functional response*, a concept initially introduced by (Solomon 1949), and later enhanced by (Holling 1959a, Holling 1959b), who introduced other types of functional response.

Other forms of interaction term, such as different positive powers of the product *N*^*k*^*P*^*m*^ other than *k* = *m* = 1, Holling’s disk equation of Type II, or sigmoid Type III functional response, are certainly possible. For example, (Braun 1993) proposed a better model for the interaction between *Paramecium caudatum* and *Didinium nasutum*, two types of single-celled protists organisms that exhibit a predatorprey relationship, by employing the following system of differential equations:

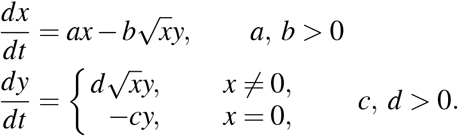

*Didinium nasutum* is a carnivorous ciliate that actively hunts and consumes *Paramecium caudatum*. See also (Chakraborty 2006) and (Harrison 1995) for other proposed models. We admit that a discussion on this topic should be addressed elsewhere.

Because there are no external factors that influence the predator and prey populations, the Lotka-Volterra model is an example of a *closed system*. This means that factors affecting the system are contained within it. On the other hand, in an *open system*, external factors can affect the dynamics of the variables. In this context, the predator and prey populations could rise or fall because of those external factors (Brown 2007).

Furthermore, although we could, we do not impose any initial conditions on the system because our purpose is not to seek a particular solution but a general solution instead. Additionally, understanding global dynamics would be more interesting than focusing on a single dynamic of the solution.

### 2.1 Exact solution and its properties

In this subsection, we use normalized variables to solve the system of ODEs. By scaling the independent variable and both dependent quantities, we reduce the number of parameters to only one. The equations become simpler, making them easier to analyze, and allowing us to focus on the underlying dynamics of the model.

By introducing the following variables:

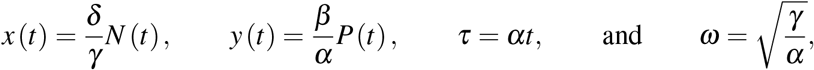

we obtain the normalized Lotka-Volterra equations:

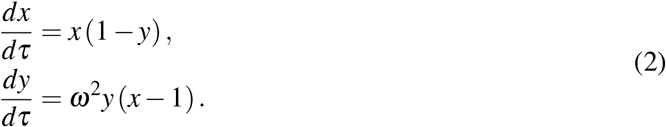

By eliminating the temporal variable *τ* from the system (2), we obtain a separable first-order ODE for the solution curves. By writing *y* = *y* (*x*) and deducing using the chain rule, the ODE is given as follows:

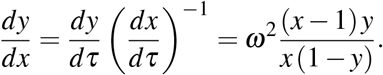

By separating the variables, that is, bringing all the *y* and *x* variables to the left and right sides of the equations, respectively, integrating both sides of the equation, we obtain a general solution of the normalized Lotka-Volterra equations:

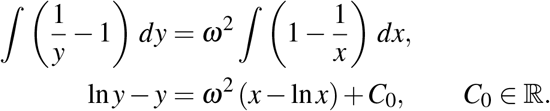

We can also express this exact solution in the following implicit form (Teschl 2012):

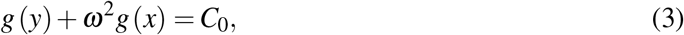

where

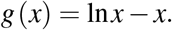

Another possible expression takes the following form (Olinick 2014):

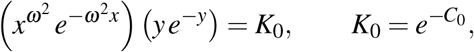

which is obtained by exponentiating each side of equation (3). In all cases, the expressions are implicit rather than explicit.

We have the following theorem regarding the solution function. This theorem confirms that the population equilibrium at (1, 1) is a local maximum (or a local minimum, depending on which two-variable function we select), implying that both populations tend to oscillate around this balanced state, a fact that might not be obvious at this stage but it will become clear as we explore the stability of equilibrium points, particularly in Subsection 2.2. Figure 2 shows three-dimensional plots of two different surfaces where critical point (1, 1) can either yield a local maximum or minimum value.

**Figure 2.**
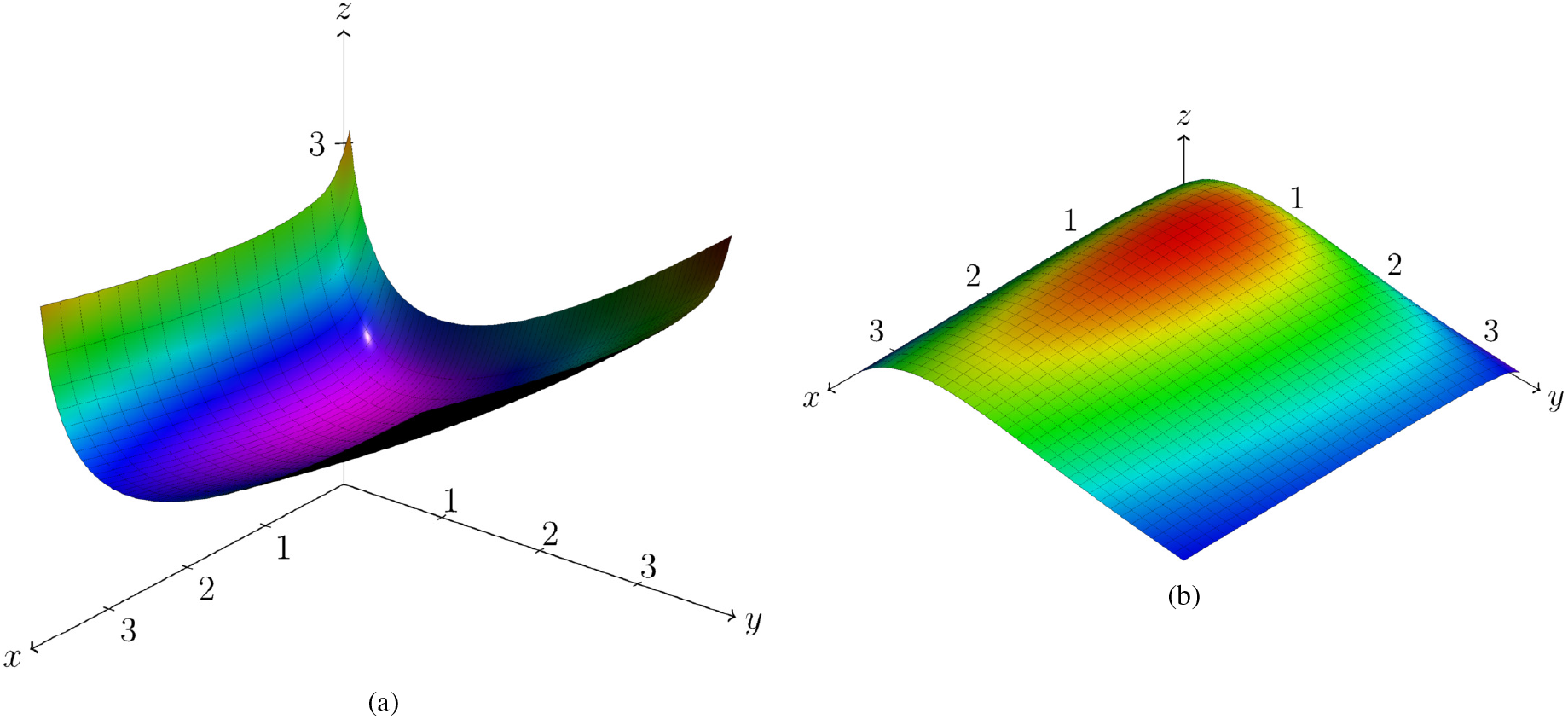
Three-dimensional renderings of the surface (a) *h* (*x, y*) = *g* (*y*) + *ω*^2^*g* (*x*) where the equilibrium point (1, 1) gives a local minimum value of *h* (1, 1) = 1 + *ω*^2^, and (b) 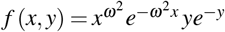 where the equilibrium point (1, 1) gives a local maximum value of 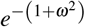. In both cases, we take *ω* = 0.5, and hence the local minimum and local maximum values are 5*/*4 and *e*^−5*/*4^, respectively.

#### Theorem 1.

*Consider a two-variable function*

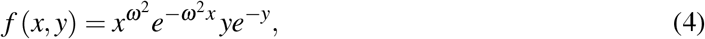

*then f* (*x, y*) *has a local maximum at* (*x, y*) = (1, 1) *with the local maximum value of* 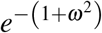.

*Proof*. The first-order partial derivatives of *f* are given by

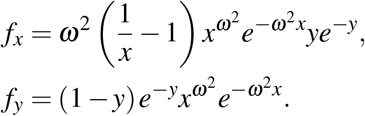

Equating both partial derivatives to zero, we obtain the critical values of *f*, i.e., (0, 0) and (1, 1).

The second-order partial derivatives of *f* are given as follows:

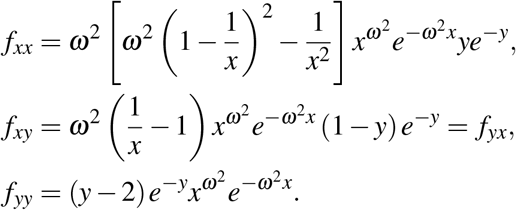

The Hessian matrix *H* of *f* is given by

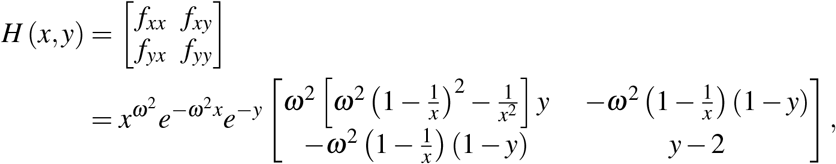

and its determinant 𝒟 is given by

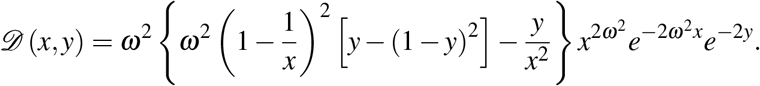

We observe that neither *H* nor 𝒟 is defined for the trivial critical point (*x, y*) = (0, 0); thus, we focus on the nontrivial critical point (*x, y*) = (1, 1). By applying the Second Derivative Test for two-variable functions, we observe that

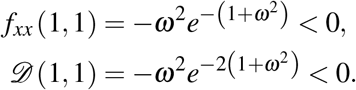

Because both values are negative, *f* has a local maximum at (*x, y*) = (1, 1) with the local maximum value of 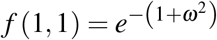.

Alternatively, we observe that the Hessian matrix at the nontrivial critical point is given by

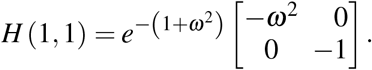

The eigenvalues of *H* (1, 1) are 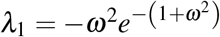 and 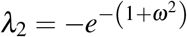. Because both eigenvalues are negative, *H* (1, 1) is a negative definite matrix. The Hessian matrix’s negative definiteness indicates that the function’s critical point is a local maximum. Thus, indeed, *f* attains an isolated local maximum at (*x, y*) = (1, 1). This completes the proof.

*Remark 1*. The choice for the two-variable function *f* (*x, y*) as in (4) is far from unique for understanding the characteristics of the exact solution of the Lotka-Volterra equation. If we choose another function, let’s say, *h* (*x, y*) = ln *f* (*x, y*), that is, the left side of equation (3), we would obtain a different Hessian matrix, its determinant, its associated eigenvalues, and its local maximum value even though the nontrivial critical point remains the same. Furthermore, we can also consider the negative of either *f* (*x, y*) or *h* (*x, y*), for which the function admits a local minimum value instead of a local maximum value. In this case, the Hessian matrix at the critical point is positive definite because its eigenvalues are both positive.

To investigate the direction of the contour plots, observe again the normalized Lotka-Volterra equations (2) and divide the population quadrant into four subquadrant regions along the lines *x* = 1 and *y* = 1, which makes the critical point as the center of these regions. See the left panel of Figure 4. Using the values of *x* and *y* in each of these subquadrants, we can investigate whether the value of 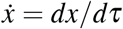 or 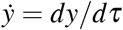 is positive or negative, and hence determine the direction of phase trajectories in each region, which is shown on the right panel of Figure 4. The result is summarized in Table 1.

**Table 1:**
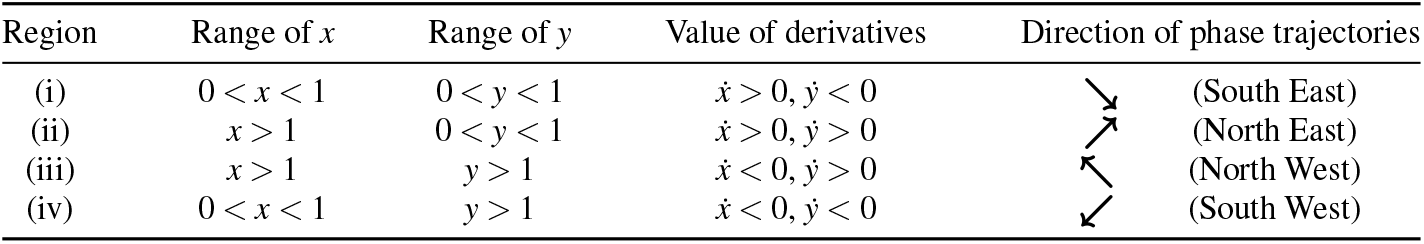
The value of derivatives 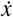 and 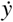 for each of the four regions depicted in Figure 4 and the direction of the phase trajectories in that particular subquadrant.

| Region | Range of $x$ | Range of $y$ | Value of derivatives | Direction of phase trajectories |
| --- | --- | --- | --- | --- |
| (i) | $0 < x < 1$ | $0 < y < 1$ | $\dot{x} > 0, \dot{y} < 0$ | (South East) |
| (ii) | $x > 1$ | $0 < y < 1$ | $\dot{x} > 0, \dot{y} > 0$ | (North East) |
| (iii) | $x > 1$ | $y > 1$ | $\dot{x} < 0, \dot{y} > 0$ | (North West) |
| (iv) | $0 < x < 1$ | $y > 1$ | $\dot{x} < 0, \dot{y} < 0$ | (South West) |

In what follows, we will demonstrate that the orbits of the solution curves of the system (2) originating in the first quadrant form a family of closed and periodic curves. The proof is available in (Braun 1993, Zill 2023a), and we include and rewrite it to enhance clarity.

#### Theorem 2.

*All solution trajectories of the system* (2) *consist of a family of closed curves in the first quadrant of the xy-plane*.

*Proof*. Let *f* in (4) be expressed as the product of separable functions, i.e., *f* (*x, y*) = *η* (*x*) *θ* (*y*) for *x* ≥ 0 and *y* ≥ 0, where

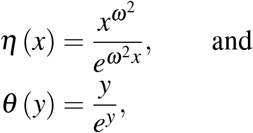

then we have the following properties:

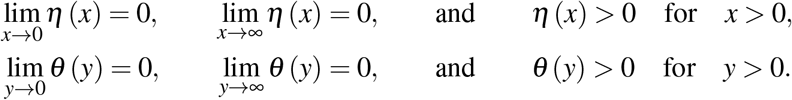

The derivatives of these two functions are given as follows:

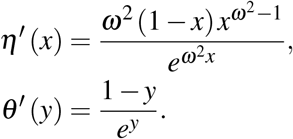

Because *x* = 1 and *y* = 1 are critical points for *η* (*x*) and *θ* (*y*), respectively, *η* (*x*) and *θ* (*y*) each attain an absolute maximum value at *x* = 1 and *y* = 1, respectively. The maximum values are given as follows:

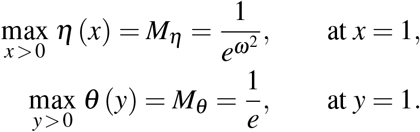

Figure 5 displays the graphs of the functions *η* (*x*) for a particular value of *ω* and *θ* (*y*), where both functions reach their maximum values at *x* = 1 and *y* = 1, respectively.

Except for the values of zero and the absolute maxima, each of the functions *η* (*x*) and *θ* (*y*) takes on all values in their range precisely twice. As a consequence, we have the following situations:

- The function *f* (*x, y*) = *K*_0_ has no real-valued solutions for *x* > 0 and *y* > 0 when *K*_0_ > *M*_*η*_ *M*_*θ*_ .
- The function *f* (*x, y*) = *K*_0_ has exactly one solution, namely (*x, y*) = (1, 1) when *K*_0_ = *M*_*η*_ *M*_*θ*_ .
- When *K*_0_ < *M*_*η*_ *M*_*θ*_, we obtain several sub-possibilities. Let us therefore consider the case where *K*_0_ = *LM*_*θ*_, where 0 < *L* < *M*_*η*_ . Observe that the function *η* (*x*) = *L* admits two solutions *x* = *x*_*m*_ < 1 and *x* = *x*_*M*_ > 1. See Figure 5(a). Thus, the critical point (*x* = 1) lies between *x*_*m*_ and *x*_*M*_, i.e., *x*_*m*_ < 1 < *x*_*M*_.
  ∘ If *x* is outside the interval [*x*_*m*_, *x*_*M*_], i.e., 0 < *x* = *x*_*k*_ < *x*_*m*_ or *x* = *x*_*K*_ > *x*_*M*_, then

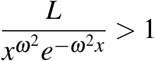

and

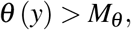

which implies that *θ* (*y*) has no real-valued solutions. See Figure 6(a).
  ∘ If *x* = *x*_*m*_ or *x* = *x*_*M*_, then

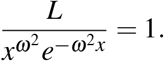

Hence, *θ* (*y*) = *M*_*θ*_ and it has a single solution *y* = 1. The red dashed lines in Figure 6(a) show these intersections.
  ∘ If *x*_*m*_ < *x* < *x*_*M*_, then

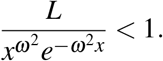

Consequently, *θ* (*y*) < *M*_*θ*_ has exactly two solutions *y*_1_ (*x*) and *y*_2_ (*x*) that satisfy 0 < *y*_1_ < 1 < *y*_2_. Additionally,

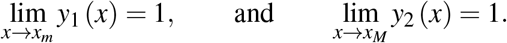

Figure 5(b) shows these solutions *y*_1_ and *y*_2_ in the graph of *θ* (*y*) and Figure 6(b) displays the corresponding values in the periodic solution in the *xy*-plane.

From this analysis, we conclude that the curves defined by *f* (*x, y*) = *K*_0_ are closed trajectories in the first quadrant of the *xy*-plane, as depicted in Figure 6. The proof is complete.

### 2.2 Population equilibrium and its stability

Population equilibrium occurs when the growth rates of both species stabilize, resulting in no net change in population levels. In this state, the predator and prey populations reach a dynamic balance. These constant solutions of an autonomous system are called *equilibrium solutions*. Mathematically, they are given as follows:

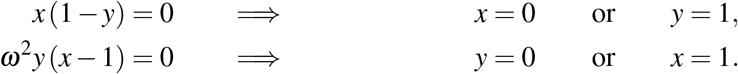

The corresponding point orbits in the *xy*-plane are called the *equilibrium points*, which are located in the *population quadrant* (that is, the first quadrant of the *xy*-plane with *x* ≥ 0 and *y* ≥ 0). It follows that there are two equilibrium points:

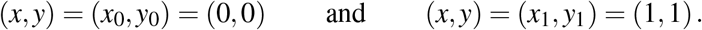

These equilibrium points are also called *critical points* or *fixed points* in the literature.

*Remark 2*. In the original variables, the nontrivial equilibrium point corresponds to

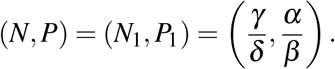

On the one hand, the first equilibrium solution effectively represents the extinction of both species. If both populations are at zero level, then they will continue to be so indefinitely. On the other hand, the second equilibrium solution represents a fixed point at which both populations sustain their current, non-zero numbers indefinitely.

To assess the stability of equilibrium points, we can analyze small perturbations around these points using the *Jacobian matrix*, which represents the sensitivity of each species’ growth rate to changes in population size. Let

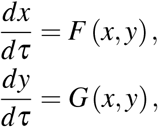

be a system of ODEs. Let (*x, y*) = (*ξ*_0_, *ζ*_0_) be an equilibrium point of the system; then by Taylor-expanding the expressions on the right-hand side about the equilibrium point, we obtain

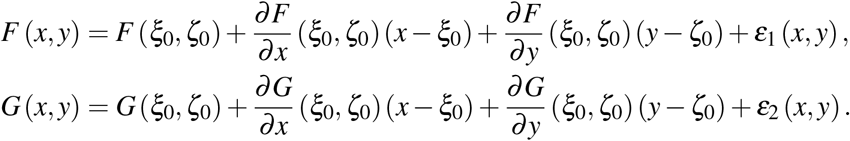

Let 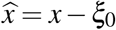 and 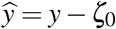, then the associated linearized system can be written as

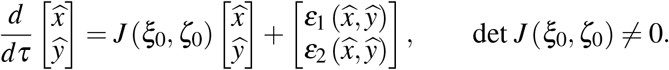

In this expression, the matrix *J* is called the *Jacobian* or *community matrix* (Kot 2001, Murray 2007), and defined as follows:

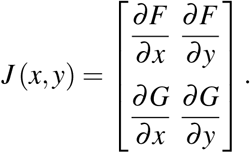

In particular, the Jacobian associated with the Lotka-Volterra model is given by

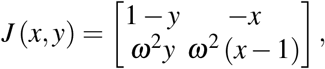

which can be easily obtained by taking partial derivatives of *F* (*x, y*) = *x* (1 − *y*) and *G* (*x, y*) = *ω*^2^*y* (*x* − 1) with respect to *x* and *y*.

The Jacobian matrix provides insight into the local behavior of the system near equilibrium points, playing an essential role in determining stability and system response to small perturbations. Having identified equilibrium points, we now explore their stability, starting with the trivial equilibrium point.

#### 2.2.1 Extinction equilibrium point

Let us consider the equilibrium point at the origin (*x*_0_, *y*_0_) = (0, 0), which is called the *extinction equilibrium point*. The associated community matrix *J*_0_ is given by

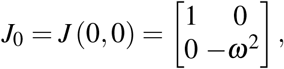

with eigenvalues

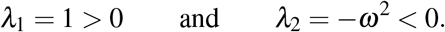

This equilibrium point is a saddle point, and hence unstable. If it were stable, nonzero populations might be attracted towards it. Such dynamics might lead towards the extinction of both species for many cases of initial population levels.

#### 2.2.2 Oscillation equilibrium point

In a predator-prey system, populations typically do not stabilize at a single value but instead oscillate around equilibrium points. Prey populations tend to rise and fall, followed by similar oscillations in predator populations, with a slight delay due to their dependency on prey availability.

In this subsubsection, we investigate the characteristics of the *oscillation equilibrium point* at (*x*_1_, *y*_1_) = (1, 1). The associated community matrix *J*_1_ is given by

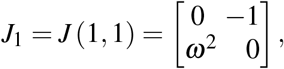

with both eigenvalues purely imaginary and conjugate to each other:

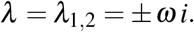

For this case, the equilibrium point must either be a center with a closed orbit or a spiral point. It turns out that (*x*_1_, *y*_1_) = (1, 1) is a stable center equilibrium point. Both predator and prey populations oscillate indefinitely around this equilibrium point.

To determine the trajectory of the closed orbit around this equilibrium point, consider again the linearized system. From

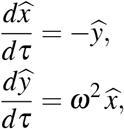

we obtain

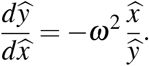

Separating the variables and integrating both sides of the equation, we obtain a family of ellipses centered at (1, 1):

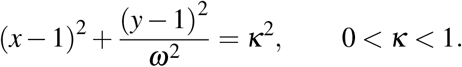

If 0 < *ω*^2^ < 1, the ellipses are elongated somewhat in the horizontal direction. Furthermore, for *κ* → 0, the ellipse shrinks to the equilibrium point (1, 1), whereas for *κ* → 1, the value of *x* may become zero, which is a contradiction to the assumption that *x* > 0.

Figure 7 shows a family of ellipses centered at the oscillation equilibrium point (*x, y*) = (1, 1) for a particular value of *ω* and several values of *κ*. It satisfies Volterra’s First Law, which states that if the system is undisturbed, the predator and prey populations would evolve and stay on their orbit. For a particular initial condition, the orbit would be traced counterclockwise. The situation also occurs for the nonlinearized Lotka-Volterra system, whose contour plots were depicted in Figure 3.

**Figure 3.**
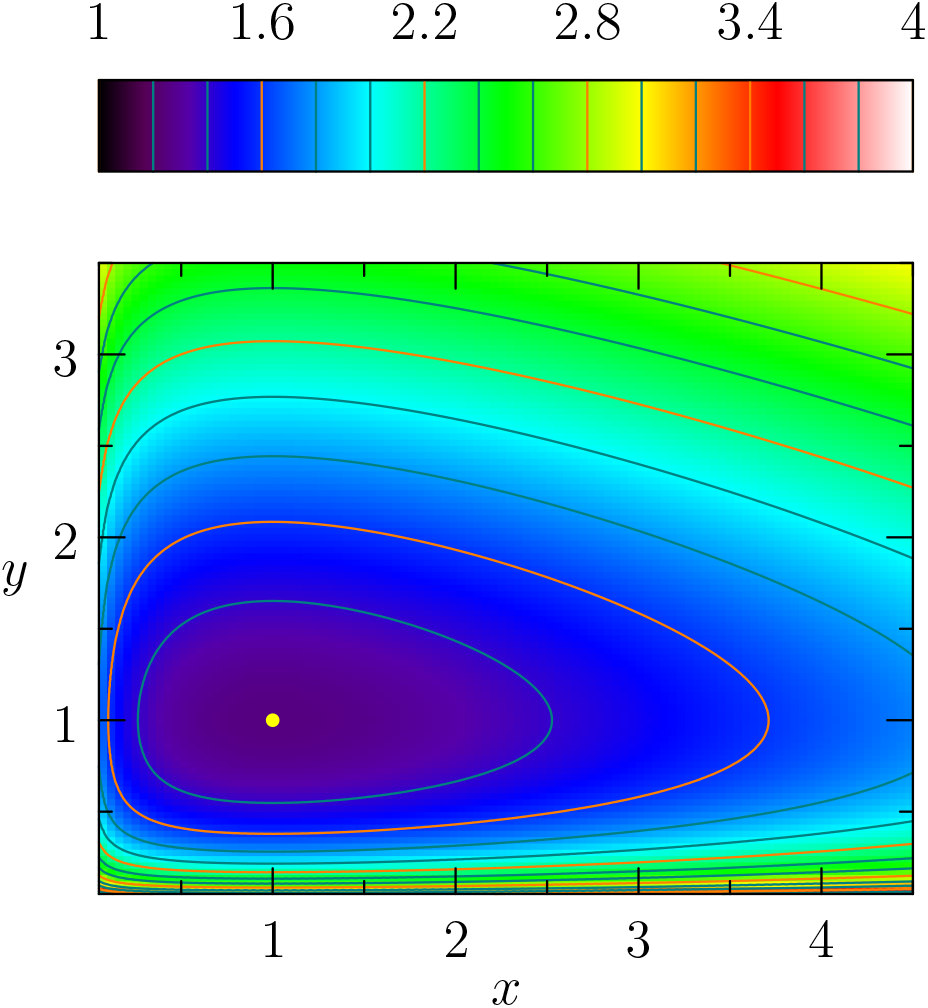
Contour plots of the surface *h* (*x, y*) = *g* (*y*) + *ω*^2^*g* (*x*) for *ω* = 0.5. The critical point (*x, y*) = (1, 1) is shown as a yellow dot.

**Figure 4.**
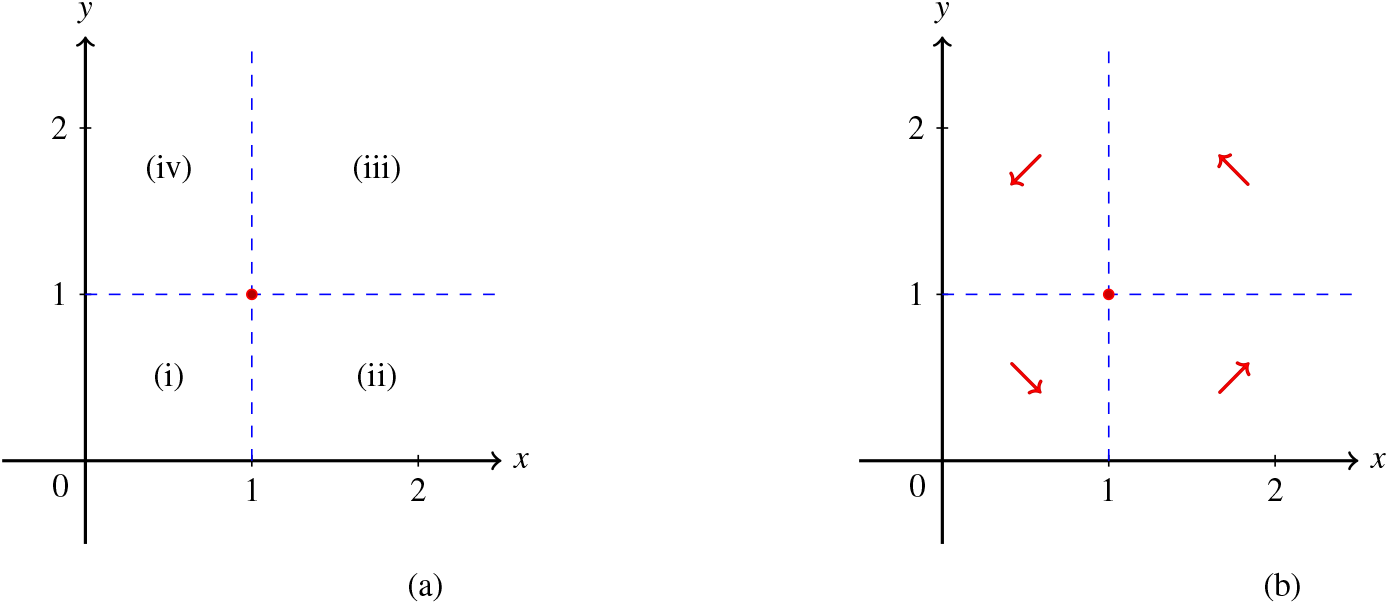
(a) The population quadrant is divided into four subquadrants with the center at (1, 1). The naming of the region proceeds in the counterclockwise direction, starting at the bottom left subquadrant and ending at the top left region. (b) The direction of phase trajectories in each of the region.

**Figure 5.**
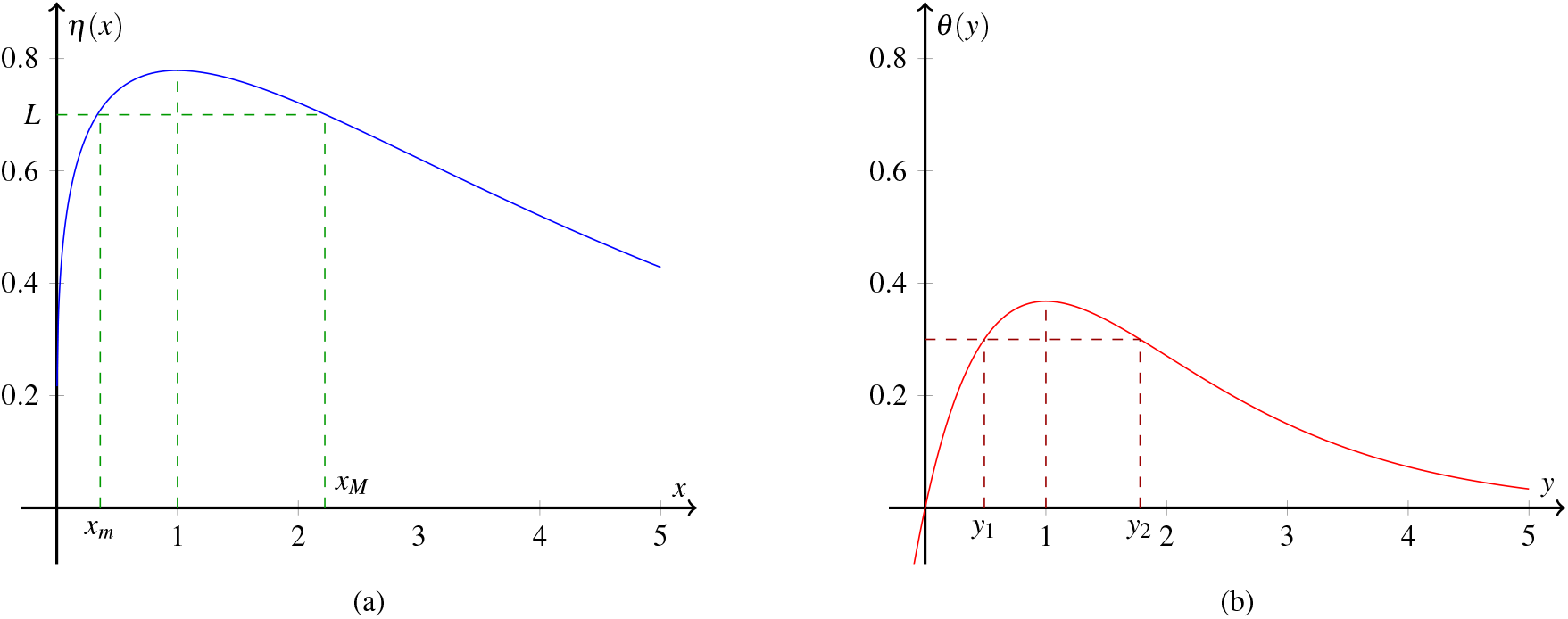
(a) Graph of the function *η* (*x*) for *ω* = 1*/*2. It reaches the absolute maximum value of *M*_*η*_ = *e*^−1*/*4^ at *x* = 1. For *L* < *M*_*η*_, the curve has two solutions *x*_*m*_ and *x*_*M*_, where *x*_*m*_ < 1 < *x*_*M*_. (b) Graph of the function *θ* (*y*), which reaches the absolute maximum value of *M*_*θ*_ = 1*/e* at *y* = 1. The solutions *y*_1_ and *y*_2_ exist when we combine both functions to form the two-variable function *f* (*x, y*) = *K*_0_ satisfying *K*_0_ < *M*_*η*_ *M*_*θ*_ and *x* ∈ [*x*_*m*_, *x*_*M*_].

**Figure 6.**
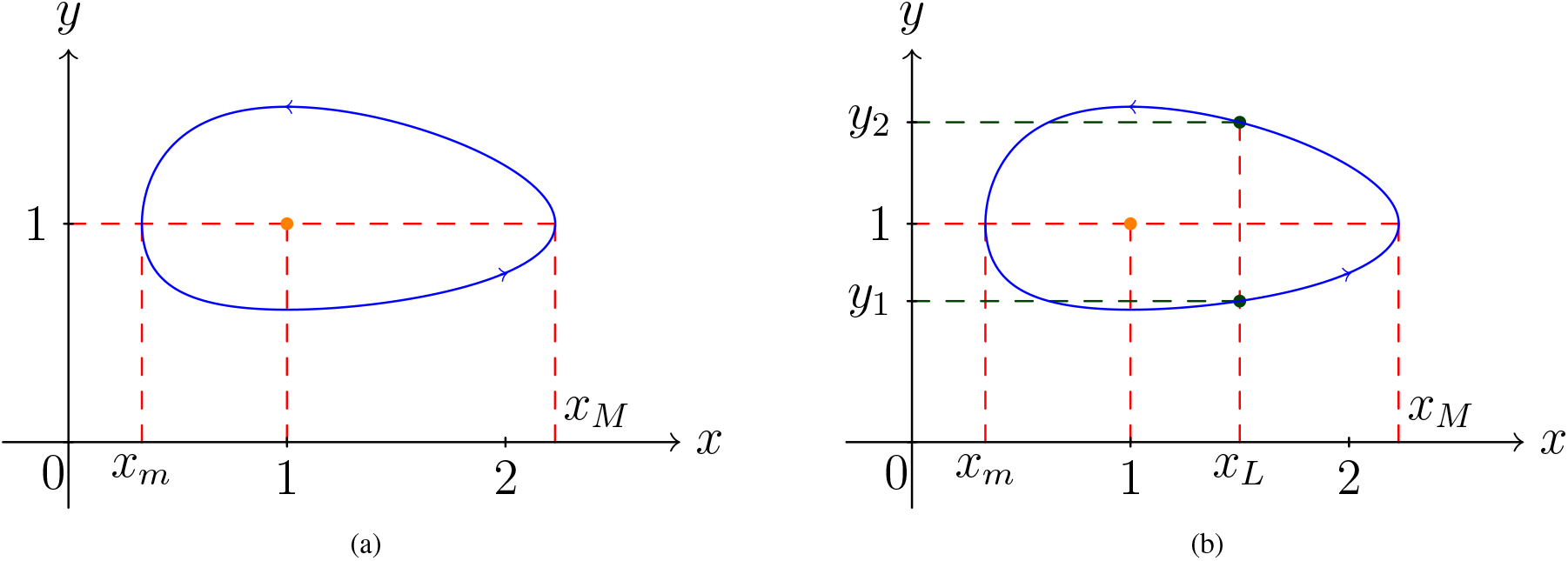
A periodic solution of the Lotka-Volterra model traversed in the counterclockwise direction. (a) For *x* ∉ [*x*_*m*_, *x*_*M*_], there exists no value of *y* that satisfies the solution. For *x* = *x*_*m*_ or *x* = *x*_*M*_, *y* = 1 is the only solution. (b) For *x*_*m*_ < *x* = *x*_*L*_ < *x*_*M*_, there are two values of *y*_1_ and *y*_2_ that satisfy the solution, where *y*_1_ < 1 < *y*_2_.

**Figure 7.**
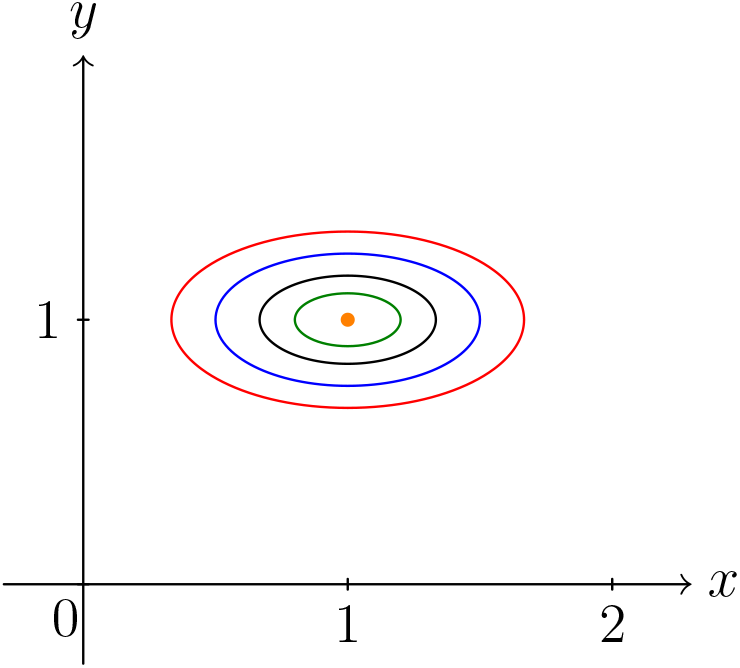
A family of ellipses centered at (*x, y*) = (1, 1) (shown as the orange dot) for *ω* = 0.5 and several values of *κ*: *κ* = 0.2 (green), 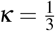 (black), *κ* = 0.5 (blue), and 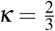 (red). The trajectories trace in the counterclockwise direction, as explained in Table 1 and Figure 4.

To express an explicit solution for the linearized system for the oscillation equilibrium point, we first need to calculate the eigenvectors. For *λ*_1_ = −*ω i* and *λ*_2_ = *ω i*, the associated eigenvectors **u**_1_ and **u**_2_ are given as follows:

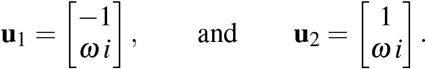

The solution to the linearized system is then given by:

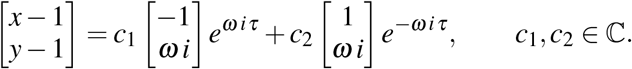

Or, by defining the complex constants *c*_1_ and *c*_2_, we can express the solutions with real-valued constants:

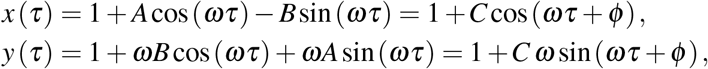

where the constant *C* and phase *φ* are determined by the initial conditions, and the following relations hold:

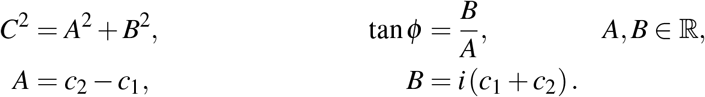

From the obtained solutions, we can observe the following characteristics:

- The average populations are 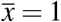 and 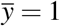. See Theorem 3.
- The sizes of the predator and prey populations vary sinusoidally with period *T* = *T*_0_ = 2*π/ω*, independent of the initial conditions.
- The predator and prey populations are out of phase by one-quarter of a cycle. The prey leads and the predator lags.

Figure 8 depicts the graphs of the solution of the linearized Lotka-Volterra system for a selected value of parameters, showing the periodic nature of both predator and prey populations.

**Figure 8.**
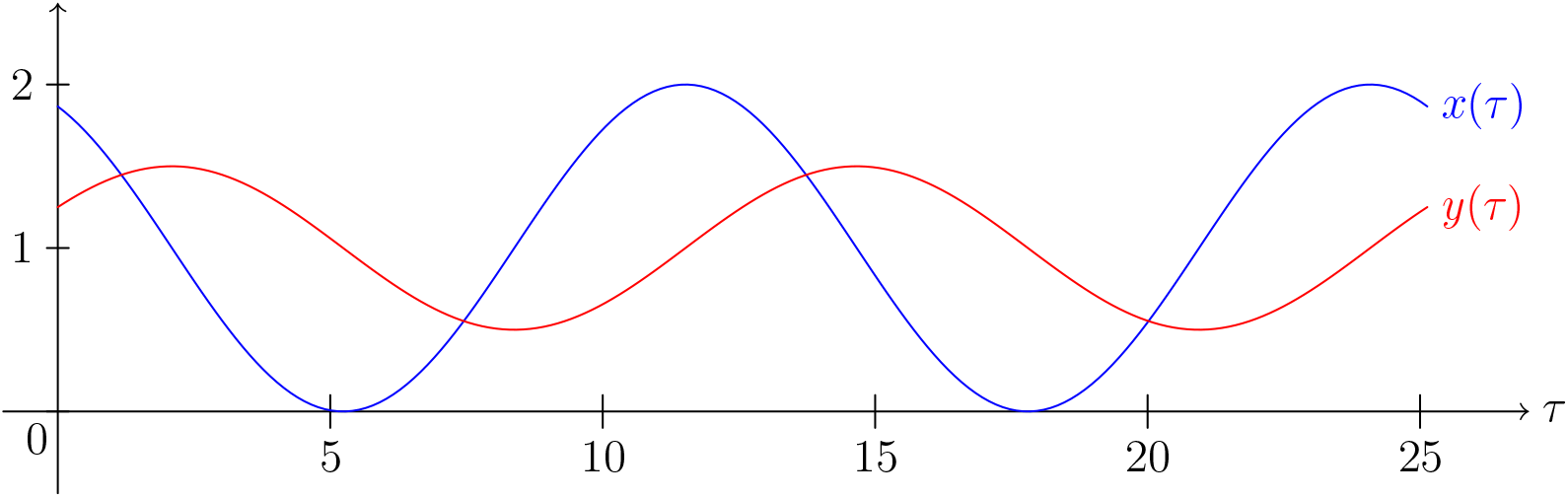
The solutions of the linearized system for *C* = 1, *ω* = 0.5, and *φ* = *π/*3. The normalized prey *x* and predator *y* variables are depicted in blue and red curves, respectively.

##### Theorem 3.

*Let x* (*τ*) *and y* (*τ*) *be a periodic solution of the Lotka-Volterra model* (2) *with period T* > 0. *The average sizes of the prey and predator population densities are defined as*

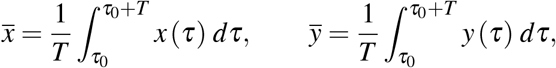

*where τ*_0_ ≥ 0. *Then, the average values of x* (*τ*) *and y* (*τ*) *are the equilibrium values, i.e*., *x* = 1 *and y* = 1.

*Proof*. Divide both sides of the second equation of (2) by *y*, integrate with respect to *τ* for one period from *τ*_0_ to *τ*_0_ + *T*, and divide the obtained expression by its period *T* to obtain:

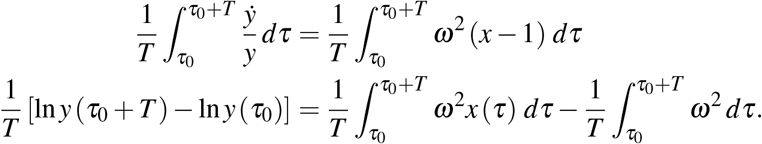

Because the function is periodic, i.e., *y* (*τ*_0_) = *y* (*τ*_0_ + *T* ), the left side of the equation vanishes. Furthermore, because *ω*^2^ > 0, we can cancel it from the right-hand side of the equation. It follows that

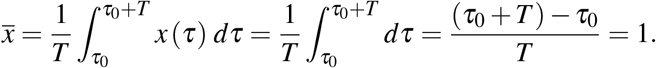

In a similar manner, we can also verify that 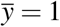. The proof is complete.

While the classical Lotka-Volterra model offers valuable insights, it has limitations. A modified version of the Lotka-Volterra model is one where the prey population follows the logistic model instead of exponential model, in addition to its loss due to kills by the predator (Baker 2016, Brown 2007, Conrad 2003, Noonburg 2014, Olinick 2014, MacCluer et al. 2019). Incorporating environmental variables such as habitat quality or climate conditions makes the model more adaptable to real-world ecosystems under stress, which is essential for sustainability planning.

### 2.3 Oscillation period

#### 2.3.1 Period calculation

Although the period of the linearized Lotka-Volterra system is easily identified, i.e., *T*_0_ = 2*π/ω*, the period of the original system cannot be expressed in terms of elementary functions. (Shih 1997) surveyed several integral representations for this period and although they look different, he verified that they were equivalent. Volterra himself derived a formula for the period and expressed it as the sum of four integrals corresponding to the four segments of the closed orbit of the system (Volterra 1926b). By employing perturbation techniques, an asymptotic formula for the period associated with each of the four integrals can further be obtained, as demonstrated by (Grasman and Veling 1973).

By reducing the normalized Lotka-Volterra system to a van der Pol-type equation and rewriting the second-order differential equation as an auxiliary system of differential equations, (Hsu 1983) expressed the period as a summation of two integrals and also showed that the period is a strictly increasing function of the energy level determined by initial values *x*_0_ and *y*_0_.

Using the theory of Hamiltonian systems, Waldvogel and Rothe also derived integral expressions for the period. The former expressed the Lotka-Volterra equations as a Hamiltonian system with one degree of freedom and using a suitable transformation, represented the period as the integral over a full period of a continuous periodic function (Waldvogel 1983, Waldvogel 1986). The latter formulated the period as a convolution integral of the Laplace transform of the energy-period function (Rothe 1985).

Using the Lindstedt-Poincaré technique, periodic solutions of the Lotka-Volterra system can be approximated and its period can be estimated accordingly (Grozdanovski and Shepherd 2008). It is given explicitly by

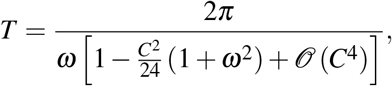

where *C* is a small amplitude that depends on the initial conditions previously derived in Subsubsection 2.2.2.

In this subsubsection, we outline the derivation by employing the Lambert *W* function, which is the inverse of the function *xe*^*x*^, following the argument presented by (Shih 1997). We define the energy level of the Lotka-Volterra system as follows.

##### Definition 1.

The energy level *E* is given by the difference between the first integral and its value at the minimum value, i.e.,

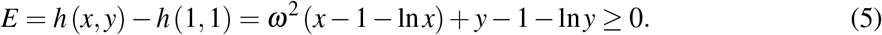

We now have the following theorem.

##### Theorem 4.

*The closed trajectory given by the energy level* (5) *admits the following period T, which depends on E:*

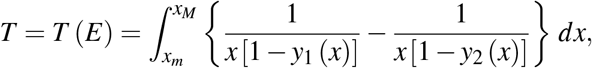

*where the functions y*_1_ *and y*_2_ *correspond to the lower and upper parts of the closed trajectory solution, respectively, and are expressed in terms of the Lambert W function, i.e*.,

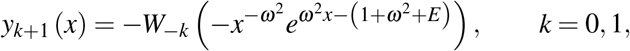

*and the lower and upper boundaries of integration x*_*m*_ *and x*_*M*_ *are the minimum and maximum values of x, respectively, and are also expressible in terms of the Lambert W function, i.e*.,

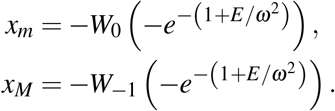

*See Figure 6 for an illustration of the periodic solution*.

*Proof*. Begin with the expression of the energy *E* (5), exponentiate it, and separate the variables:

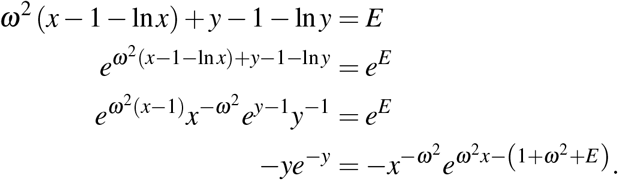

This equation admits the following solutions in terms of the Lambert function *W* :

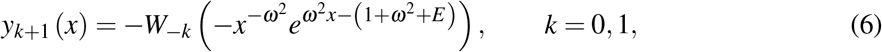

for *y* > 0 and

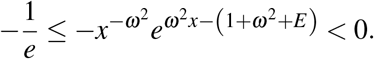

By canceling out the factor *e*^−1^, bringing the variable terms to the left-hand side, and taking the entire expression to the power of 1*/ω*^2^, we obtain the following inequality:

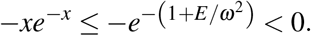

The range for *x* that satisfies this inequality is between the minimum value *x*_*m*_ and the maximum value *x*_*M*_, where

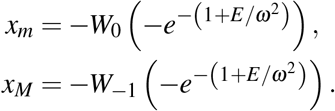

Substituting the solutions (6) into the first equation of the normalized Lotka-Volterra equation (2) and separating the variables yields

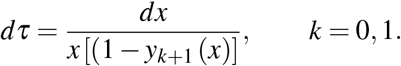

Traversing along the lower portion of the solution trajectory *y*_1_ (*x*) in the counterclockwise direction from the point (*x*_*m*_, 1) on the left with *τ* = *τ*_*m*_ to the point (*x*_*M*_, 1) on the right with *τ* = *τ*_*M*_, we obtain the following integral:

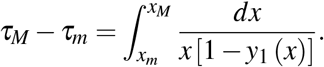

Similarly, by traversing along the upper part of the solution trajectory *y*_2_ (*x*) in the counterclockwise direction from the point (*x*_*M*_, 1) on the right with *τ* = *τ*_*M*_ to the point (*x*_*m*_, 1) on the left with *τ* = *τ*_*m*_, it yields the following expression:

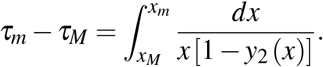

By combining both expressions for the integral, we obtain the integral representation of the period *T* . The proof is complete.

##### Corollary 1.

*Let A*_*x*_ *and A*_*y*_ *be the amplitudes for the prey and predator population densities, respectively, then:*

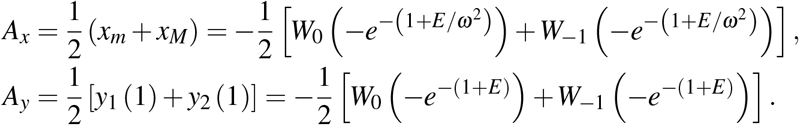

The proof can be worked out using the definition of the amplitude, i.e., half the vertical distance between a crest and a trough. For the prey solution *x*, the crest and trough occur at *x*_*M*_ and *x*_*m*_, respectively, when *y* = 1. For the predator solution *y*, the crest and trough occur at *y*_2_ and *y*_1_, respectively, when *x* = 1.

*Remark 3*. Figure 6 is produced by sketching the two-variable function *f* (*x, y*) = *K*_0_ while the period calculation employs the energy function (5). Although the two functions are quantitatively distinct, they are qualitatively identical. By selecting appropriate constant level curve *E* = *E*_0_ that matches with *K*_0_, we can obtain an identical quantitatively periodic solution depicted in Figure 6, i.e., *E*_0_ = 0.75 − ln *K*_0_. In our particular example, because we selected *K*_0_ = 0.7*/e*, we should choose *E*_0_ = −(0.25 + ln 0.7) to produce an identical trajectory in the *xy*-plane shown in Figure 6.

Using the well-known data from the Hudson Bay company on lynx and hare pelts from 1900 to 1920, we can observe that the common period *T*_*D*_ is around 10.5 years. By finding the best-fitting the predator-prey model to the parameters obtained from the data, we can calculate the common period for both the linearized system *T*_0_ as well as its improved approximation *T*_*M*_. They are given as follows, respectively:

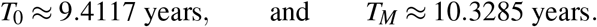

We observe that the common period obtained from fitting the model to data parameters shows better agreement with the common period found in nature, slightly differs within a couple of months, i.e., around two months only.

Additionally, we can also extract information regarding the amplitudes for both the predator and prey populations. According to the pelt data, the amplitude for hare population is around 47,750 and the lynx population is around 33,400. Our best-fit model suggests that the hare population amplitude is around 43,043 and the lynx population amplitude is around 29,708. The average difference in both population amplitudes is around 10.46%, i.e., 9.86% and 11.05% of the amplitude differences for hare and lynx populations, respectively.

By comparing the quantities obtained from the data and the best-fitting model, one can observe some intriguing similarities and differences, including the initial conditions, equilibrium point, and coefficients of the Lotka-Volterra model, as displayed in Table 2. Figure 9 displays further comparisons between the data from hare and lynx pelts and the numerical simulation by fitting the model with the best parameters. Both the evolution of each species’ population and the phase plane trajectory are shown in the left and right panels, respectively.

**Table 2:**
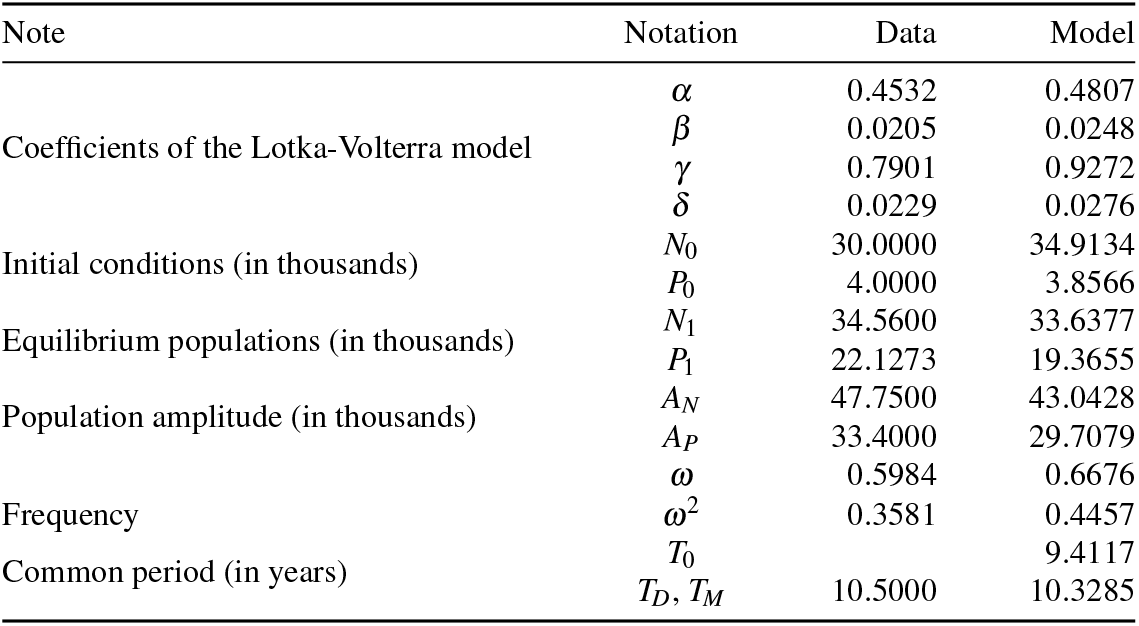
Figure comparison between the hare *N* and lynx *P* pelt data from the Hudson Bay company and the best fitting model, by minimizing the sum of square errors of the two datasets.

**Figure 9.**
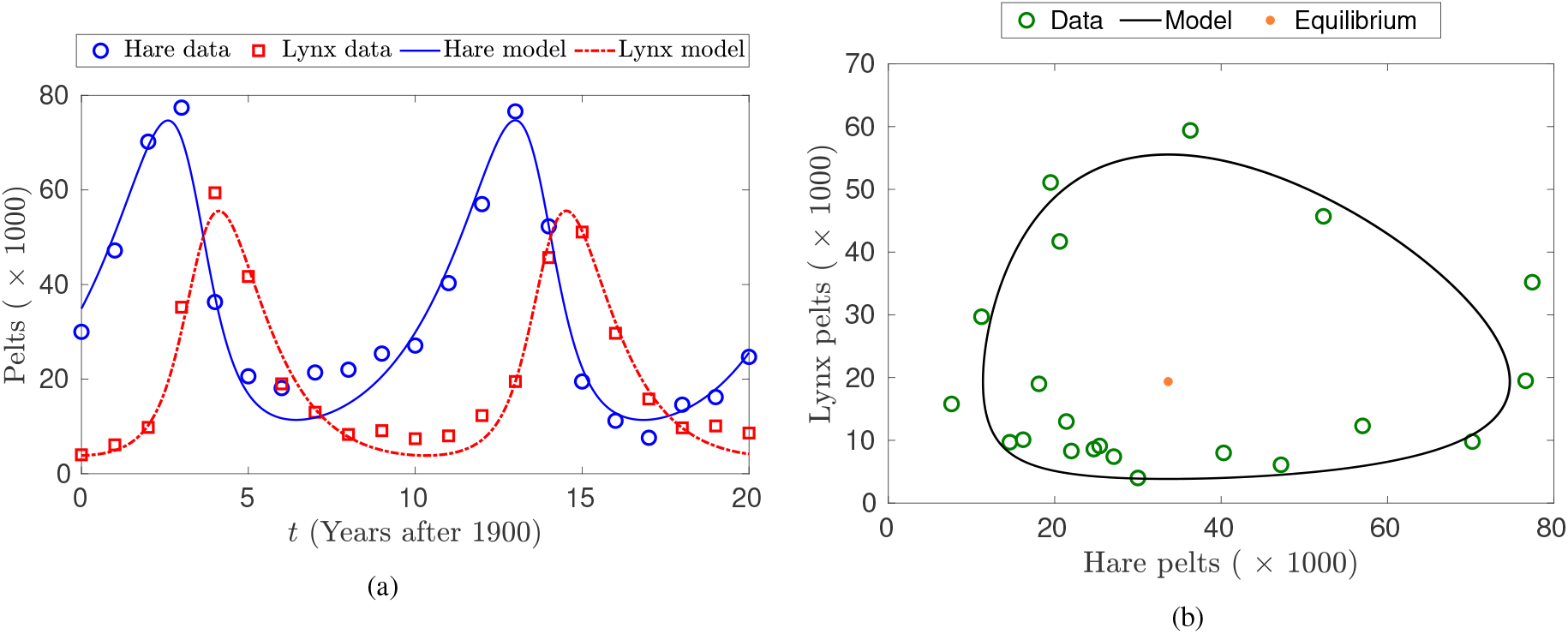
Comparison between pelt data and the Lotka-Volterra model based on best-fitting parameters shows a remarkable qualitative agreement. (a) The comparison of the time evolution for 20 years from 1900 to 1920. The hare and lynx population data are indicated by the blue circles and red squares, respectively, whereas the numerical simulation results for the hare and lynx populations are depicted by the solid blue line and red dotted-dashed curve, respectively. (b) The phase plane comparison between the pelt data and the best-fitting computation. The phase plane data are shown in scattered green circles and the predator-prey model is shown as a solid black closed curve. The equilibrium population based on the simulation is shown as the filled orange circle.

#### 2.3.2 Time scale and oscillation period

Although the time variable is normalized in this section, practitioners such as ecologists and conservationists are more interested in the real time scale of the Lotka-Volterra model. The period of predator-prey oscillations depends on multiple factors, including biological and ecological. These include the generation time and reproductive rates of both species, maturation time to reproductive age, prey recovery rates after predation pressure, environmental carrying capacity, resource availability, seasonal effects, and environmental variability (Boyce et al. 1999, Varpe 2017, Wilbur et al. 1974).

This explains why there is no unique time scale for the model as different pairs of species exhibit distinct time scales. It should not be surprising, however, that the oscillation period would be shorter than the typical lifetime of the interacting species (Lipowski and Droz 2006). For example, the oscillation period of small rodents, such as voles and lemmings, with their predators, such as owls and weasels is around three to five years (Hansson and Henttonen 1989, Hörnfeldt 1991, Korpimäki et al. 2025).

One of the most well-documented examples is the classic population interaction between lynx (*Lynx canadensis*) and snowshoe hares (*Lepus americanus*) which live in Canadian boreal forests, as we discussed comparatively in Subsubsection 2.3.1. Being relatively isolated from external confounding factors, they exhibit an approximately 10-year cycle (Elton and Nicholson 1942, Krebs et al. 2001). Other predator-prey interactions might exhibit shorter or longer cycles, depending on the species and their environment.

There are several approaches to estimating and forecasting predator-prey cycles from data. For instance, using a model-based approach, the parameters in the Lotka-Volterra model can be estimated using least-squares technique or maximum likelihood estimation (Iannelli and Pugliese 2015, Lee and Foy 2025). Other methods can be employed by using time series analysis. The method of spectral analysis identifies dominant frequencies by decomposing the time series data into constituent frequencies (Gonzalez and Loreau 2009, Saeedian et al. 2022). Furthermore, autocorrelation analysis examines how data points correlate with themselves at different time lags. Strong correlation at specific lags indicates cyclical behavior (Dray et al. 2010, Lichstein et al. 2002).

## 3 The Lotka-Volterra model and (un)sustainability

In this section, we explore the relationship between the dynamics of predator-prey interaction and sustainability. On the one extreme, Subsection 3.1 argues that a solid understanding of quantitative aspects of the model will be essential for sustainable population management. On the other extreme, Subsection 3.2 demonstrates that as countries beef up their military strength, the race to accumulate weapons can lead to conflict and war, which is unsustainable and has a significant negative impact on the environment and human beings.

### 3.1 Sustainable population management

Although the Lotka-Volterra model discussed in this chapter is the simplest model for predator-prey dynamics, it can nevertheless still can provide insight into managing populations sustainably by understanding how the interacting populations fluctuate over time. Although population management was originally concerned with societal planning and resource allocation for human populations, the principle can also be applied to ecology with its interacting species of flora and fauna.

According to the United Nations Population Fund, sustainable population management refers to strategies and practices that aim to balance population size with the availability of resources and the capacity of the environment to support life, ensuring a healthy planet for present and future generations. It is not just about limiting population growth, but also about managing resources, consumption patterns, and environmental impact (Albrectsen 2013).

To sustainably manage predator-prey populations within a given habitat, it is essential to understand their natural cyclical dynamics. Our goal should be to balance these population fluctuations to prevent overexploitation and ensure the long-term stability of the ecosystem (Sutherland 2001). Rather than attempting to maintain artificially stable populations, a key principle of sustainable management is to work with these natural cycles. As argued by (Grafton and Silvia-Echenique 1997), in addition to using theoretical models and simulations, wildlife managers should comprehend ecosystems and their extensive relationships among species.

This approach not only fosters more resilient ecosystems but also ensures more sustainable resource use over the long term. Achieving this objective involves various strategies, such as habitat management, carefully controlling predator populations, and setting sustainable harvest levels for both predator and prey species. For example, if the intrinsic growth rate is 0.8 per year, then harvest rates must be significantly below this value to ensure sustainability (Lande et al. 1997).

Indeed, understanding cycle patterns enables conservation managers and wildlife biologists to predict when predator and prey populations will reach their peaks and troughs (Myers 2018). In turn, this oscillation pattern could be identified as the optimal time frame for sustainable harvesting. On the one hand, harvesting prey species during the ascending phase of their cycle, that is, before the peak, would maximize yield while allowing recovery. On the other hand, harvesting during the descending phase would risk accelerating population collapse (Liz 2017, Samhouri et al. 2017).

Furthermore, in terms of monitoring frequency and management planning, the knowledge of the cycle period will ensure effective management of predator and prey populations sustainably. For example, for a 10-year cycle, annual monitoring might be sufficient. For shorter cycles, such as rodents and their predators, more frequent monitoring is needed to track population changes more accurately. A decade-long cycle period would also mean spending several decades to evaluate the success of their management intervention because short-term assessments can be misleading, especially when not even one full-cycle was captured.

Returning to our discussion about harvesting, the body of published literature is ample with studies on this topic because its effect alters the interaction dynamics between the two species and mathematically, it influences the equilibrium point and its stability. At this point, we are interested in ensuring that any harvest activities would ensure the sustainability of both species in the predator-prey interaction (see Figure 10).

**Figure 10.**
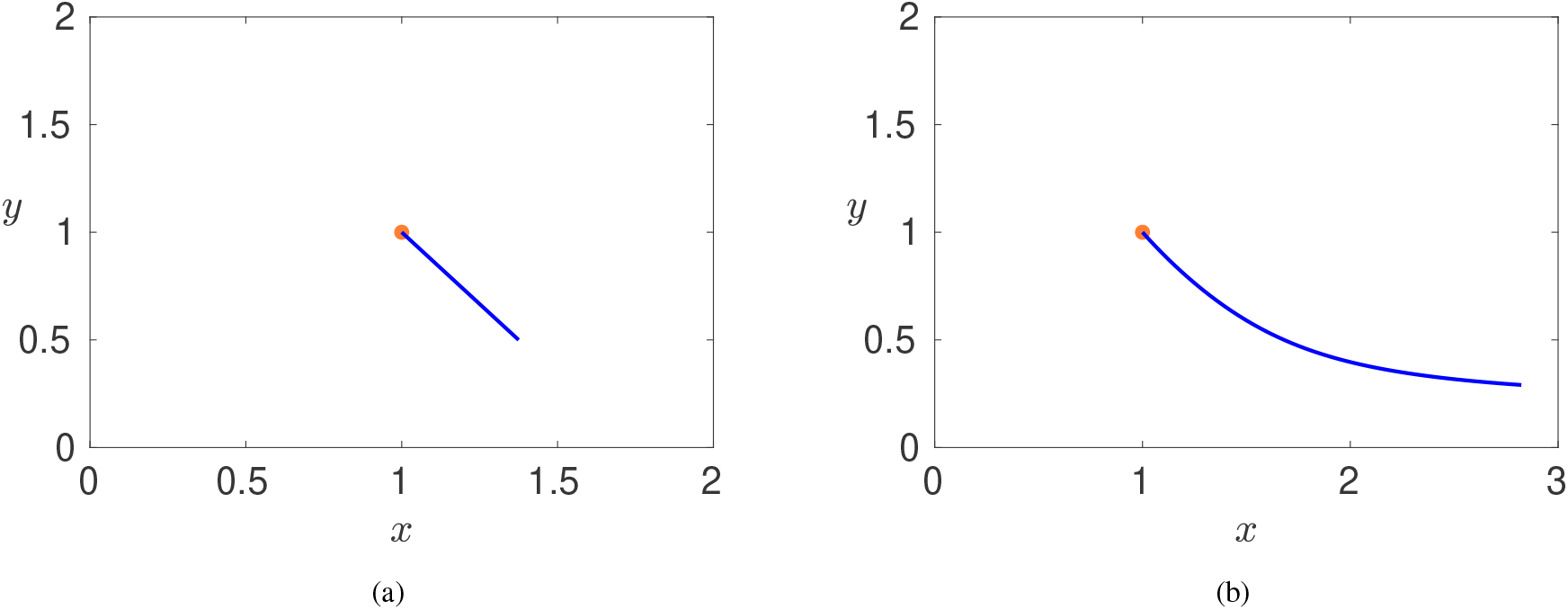
The changes in equilibrium point for a modified Lotka-Volterra model with harvesting effects. The original equilibrium point is depicted by an orange dot in both panels. (a) Linear harvesting with 0 ≤ *ε*_1_ ≤ 0.5 and *ε*_2_ = 0.75*ε*_1_. (b) Constant harvesting with 0 ≤ *H*_1_ ≤ 0.5 and *H*_2_ = 0.75*H*_1_.

One way to sustainably manage both the predator and prey species is by harvesting them in a relatively small numbers proportional to the population rate of each species, as exemplified in the case of fishing smaller food fish and larger selachian fish (Braun 1993). A mathematical model of the Lotka-Volterra equations with this kind of linear harvesting rate can be formulated by simply adding or subtracting small linear terms to the original model (2). With small positive values for *ε*_1_ and *ε*_2_ within the same order of magnitude, i.e., 0 < *ε*_1_ ≪ 1, 0 < *ε*_2_ ≪ 1, and *ε*_1_*/ε*_2_ = 𝒪 (1), the modified equations are now given as follows:

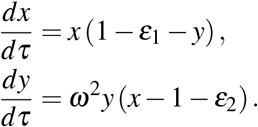

This modified system is still similar to the original model (2) but with shifted equilibrium point and also shifted average values of the predator and prey population densities, which are given by

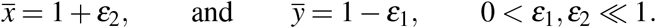

On the one hand, this implies that on average, sustainable harvesting for both species will increase the prey population density and decrease the predator’s population density. This principle is formulated as *Volterra’s Third Law*, also known as *The Law of Harvesting*, which states that constant-effort harvesting raises the average prey population density per cycle and lowers the average predator population density (Borrelli and Coleman 2004).

On the other hand, by allowing anti-harvesting, or stocking, to occur, the predator’s population will start to increase and the density of the prey population will eventually decrease because now there would be more selachians hunting those smaller fish for their meals. Mathematically, this happens when we allow a negative harvesting rate, i.e., both *ε*_1_ and *ε*_2_ become negative but still small number in their absolute sense. Thus, some good intentions and interventions that are thought to be enhancing sustainable population management turn out to yield the opposite effect.

Figure 10(a) shows how the equilibrium point evolves as we increase the value of *ε*_1_ from 0 to 0.5 and *ε*_2_ is taken to be 0.75*ε*_1_. We observe that the direction of the changes is into the fourth subquadrant from the original equilibrium point (1, 1), i.e., between the eastern and southern directions from (1, 1). For linear stocking, we would observe the evolution in the opposite direction.

Another type of harvesting that is relatively straightforward mathematically and yet our simple model might not completely capture the real situations in nature due to its limitations is the constant harvesting rate instead of linearly proportional to each population species as discussed earlier.

One of the early references of the constant harvesting is (Brauer and Sánchez 1975), who observed that there exists an upper bound such that the asymptotic stability of the equilibrium points is preserved. By analyzing the nature of the qualitative behavior of the solutions, we can comprehend and anticipate some unsustainable scenarios in real-life harvesting, including scenarios where the harvesting rate leads to the absence of equilibrium, thereby guaranteeing the predator’s extinction, or where it results in an extremely low level of population density that could be easily extinguished by a small perturbation (Brauer et al. 1976). A theoretical model was further developed to determine the region of asymptotic stability, both for asymptotically stable equilibrium points and for limit cycles of periodic solutions (Brauer and Soudack 1979a, Brauder and Soudack 1979b).

In the case of positive constant harvesting, this modified system exhibits a finite-time extinction and pattern formation, while the latter was notably absent in the case of negative harvesting, i.e., stocking, or *planting*, as the authors called it (Choi and Kim 2018). See also (Luo and Zhao 2017, Xiao and Jennings 2005) for stability and bifurcation analysis of the predator-prey systems with constant harvesting rate.

By allowing negative harvesting, i.e., stocking, we observe intriguing ecological phenomena. On the one hand, predator stocking tends not only to promote prey extinction but also to make its population dependent on the stocking. On the other hand, prey stocking tends to increase the predator population and stabilize the system (Brauer and Soudack 1981, Brauer and Soudack 1982). This finding highlights how mathematical modeling can reveal counterintuitive ecological relationships that are essential for sustainable management decisions.

Let us briefly investigate mathematically what the consequences are for the equilibrium point when we include the effect of constant harvesting *H*_1_ > 0 and *H*_2_ > 0. We modify again the original system of the normalized Lotka-Volterra model (2):

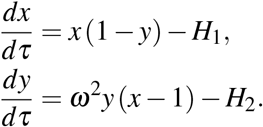

The new equilibrium points are given by (*x*_2_, *y*_2_) and (*x*_3_, *y*_3_), where

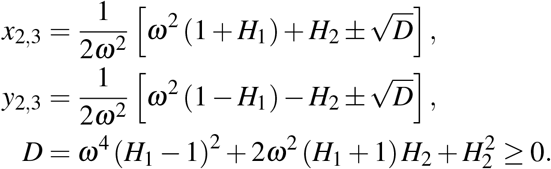

We also observe that when the harvest constants diminish, both equilibrium points return to the nontrivial and trivial equilibria, respectively, i.e.,

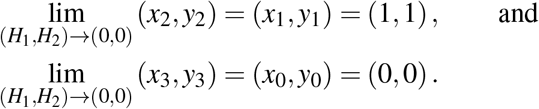

For many values of harvest constants *H*_1_ and *H*_2_, the equilibrium point related to the trivial equilibrium, i.e., (*x*_3_, *y*_3_), can be displaced outside the population quadrant and thus it does not have any meaningful ecological consequence although it is valid mathematically. Hence, as our previous interest was in the nontrivial equilibrium point, our interest here is also in the equilibrium related to it, i.e., (*x*_2_, *y*_2_).

Figure 10(b) displays the evolution of the equilibrium point under the influence of constant harvesting effects. The value of *H*_1_ ranges from 0 to 0.5 and *H*_2_ is fixed at a constant ratio with *H*_1_, in this case, *H*_2_ = 0.75*H*_1_. Similar to the case for linear harvesting effects, the equilibrium point moves in the direction of the fourth subquadrant from the original equilibrium point of (1, 1). The difference is that instead of a straight linear curve, the constant harvesting effect gives a nonlinear curve in the southeastern direction, where the rate of increase in the prey population density seems to be faster than the decrease in the predator’s population density.

We have the following theorem.

#### Theorem 5.

*For constant harvesting effects, the associated equilibrium in the population quadrant is an unstable node*.

*Proof*. The equilibrium point located in the population quadrant is (*x*_2_, *y*_2_), where the characteristic equation of the Jacobian matrix at this equilibrium point is given by:

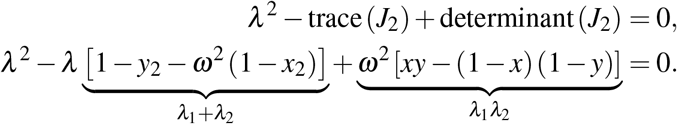

Here, *J*_2_ refers to the Jacobian matrix of the modified Lotka-Volterra system evaluated at (*x*_2_, *y*_2_). We also know that the trace of *J*_2_ is the sum of the eigenvalues and its determinant is the product of the eigenvalues. To show that (*x*_2_, *y*_2_) is an unstable node, we need to verify that both trace(*J*_2_) and determinant(*J*_2_) are positive for *ω* > 0. The latter is affirmative because the last term of the characteristic equation simplifies to 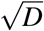, which is positive because the discriminant *D* itself is also positive. Thus, *λ*_1_*λ*_2_ > 0.

We now need to verify whether the trace is positive. Consider its product with 2*ω*^2^, which can be expressed as follows:

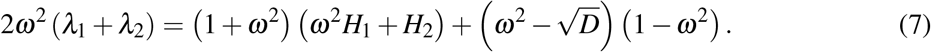

We consider two different cases:

- *ω* ≥ 1

We observe that the product of the first two terms in the parenthesis in (7) is positive, i.e.,

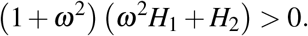

The product of the final two terms in the parenthesis in (7) is also positive because both of them are negative, i.e.,

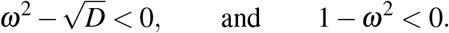

The former can be easily seen by comparing that *ω*^4^ < *D*, which implies 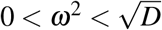. The sum of two positive terms is also positive and because the factor 2*ω*^2^ is also positive, the trace of the Jacobian *J*_2_ is also positive, i.e., *λ*_1_ + *λ*_2_ > 0.

- 0 ≤ *ω* < 1

Observe that the discriminant *D* satisfies the following inequality:

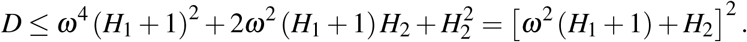

This implies

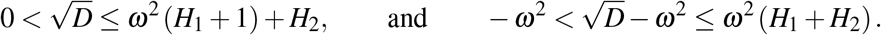

The product of 2*ω*^2^ with the trace of the Jacobian *J*_2_ (7) satisfies the following relationship:

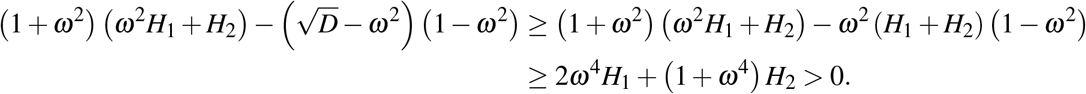

Hence, trace(*J*_2_) is positive, i.e., *λ*_1_ + *λ*_2_ > 0.

Because both the sum and the product of eigenvalues of the Jacobian *J*_2_ are positive, the equilibrium point (*x*_2_, *y*_2_) is an unstable node. This completes the proof.

This brief exposition reveals the mathematical richness of including harvesting effects in population models, a topic we barely scratch the surface of in this chapter. We observe that harvesting inherently alters the system’s equilibrium point, introducing a critical trade-off. When effectively managed, modest harvesting can be a tool for sustainable resource management, maintaining desired predator and prey populations and ensuring long-term ecosystem viability. However, if harvesting rates are too large, they can destabilize a previously stable equilibrium, potentially leading to the extinction of one or both species. Therefore, understanding these complex dynamics is crucial, as poorly managed harvesting inevitably leads to population collapses and ecological damage, undermining potential economic benefits.

### 3.2 Negative sustainability: Modified arms race model

Although the Lotka-Volterra model discussed in this chapter is the simplest model for predator-prey dynamics, its application extends beyond the realm of mathematical biology. This model is not only popular for describing ecosystems but also proves highly valuable in analyzing broad competitive phenomena across political, social, and business fields, where the nature of interactions can be effectively modeled by the Lotka-Volterra framework (Avetisyan 2024, Marino et al. 2023, Kawira et al. 2020). This broader utility encourages a deeper reflection on how to achieve balance in various competitive scenarios and contribute to sustainable development.

This subsection explores “negative sustainability,” a concept where inherent system dynamics or reactive behaviors lead to outcomes that are detrimental in the long term, consuming resources unsustainably, increasing risk, or causing collapse. The Lotka-Volterra framework, while biological, helps understand competitive interactions which, when taken to an extreme, could lead to negative sustainability.

Among the most striking historical manifestations of unsustainable competitive phenomena are wars and conflicts. These frictions have a profound negative impact on sustainability, as the destruction caused by war directly undermines efforts to achieve sustainable development goals by damaging infrastructure, displacing populations, and depleting natural resources. An important question that we should critically examine is whether mathematics and its models can contribute to preventing conflict and advocating peace, and hence promoting sustainability.

One of the mathematical models that was developed to explain the dynamics of arms races is the Richardson arms race model. Under the assumption of *mutual fear* between two nations, the arms race model describes the temporal evolution of weapons acquisition between countries (Richardson 1960). On the softer and less horrific realm, these “arms” do not have to be weapons. This pioneering work and its enduring influence in social sciences, particularly in peace and conflict studies, has been extensively documented and analyzed (Gleditsch 2020).

The classical Richardson arms race model is given by the following system of linear differential equations:

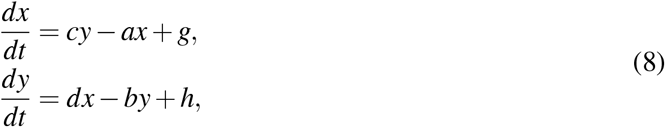

where *x* and *y* represent the arms levels of two nations, *a* and *b* represent the self-damping coefficients, that is, how much a nation’s own spending restrains further increases, *c* and *d* represent the reaction coefficients, that is, how much one nation reacts to the other’s spending, and *g* and *h* represent constant terms representing external factors of grievance and ambition. All of these coefficients usually take positive values, that is, *a, b, c, d, g*, and *h* > 0. On rare occasions, *g* and *h* can be nonpositive, that is, *g* and *h* ≤ 0, where both parties exhibit either the absence of ambition or a natural tendency toward isolation and disarmament.

For instance, let us return briefly to the field of mathematical, and particularly, evolutionary biology, where two parties are involved in adaptations and counteradaptations (McLean et al. 2024). In their classical work, British zoologists Dawkins and Krebs (1979) discussed the concept of an “arms race” in evolutionary biology, where adaptations in one lineage lead to counter-adaptations in another, potentially resulting in an unstable escalation (Dawkins and Krebs 1979).

One particular example is the relationship between cuckoo birds and their hosts. Paradoxically, cuckoo hosts are very good at detecting cuckoo eggs but not cuckoo nestlings. This suggests that cuckoos employ mimetic deception at the egg stage and direct manipulation at the nestling stage (Langmore et al. 2003, Robert and Sorci 1999, Servedio and Lande 2003). Other examples of interspecific arms races involving predators and prey have been discussed in the literature on evolutionary biology (Barber and Kawahara 2013, Hague et al. 2020, Siepielski and Beaulieu 2017).

Many of the readers of this chapter are members of a higher education institution, and some of us often wonder why such a large endowment fund is channeled toward non-academic amenities instead of improving the welfare of students and faculty. It turns out that in order to make institutions more appealing to prospective applicants, numerous colleges and universities are heavily engaged in “amenities arms races”.

In the context of higher education, the term refers to the escalating competition among various institutions to offer increasingly lavish real estate development and appealing amenities to attract “customers” but bears little relationship to formal education (Craig 2015, Hacker and Dreifus 2010, McClure 2019). In the model (8), the variables *x* and *y* refer to the level of amenity spending or luxury of two institutions, the parameters *a* and *b* mean how much a university’s own spending on amenities could internally dampens further increases, e.g., due to budget limits or donor fatigue, the parameters *c* and *d* represent how much one university reacts to another institution’s amenity spending, and the parameters *g* and *h* denote inherent desire for prestige, attracting students regardless of competitors.

Some examples include luxurious student dormitories, upscale dining halls with diverse cuisine options, entertainment venues, and extensive recreation facilities that feature lazy rivers and climbing walls. Both private and public universities, particularly in the United States, have been investing in this competition, such as High Point University in North Carolina, Texas Tech University, The University of Iowa, and California State Universities at Fullerton and Long Beach.

In the long term, this situation is not sustainable as higher education is at a crossroad, reevaluating its purpose. If colleges and universities do not build such extravagant facilities, they risk losing students to other institutions. If they do, it hurts students and parents alike who need to pay even higher costs of college enrollment. In the long term, this unsustainable situation will do more self-harm than foster flourishing for higher education institutions not only in the United States but also in other parts of the world (Varga and Lingrell 2018).

The system (8) admits a unique equilibrium point 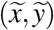 if *ab* ≠ *cd*, which is an intersection of the two nullclines *ax* − *cy* = *g* and −*dx* + *by* = *h*, where

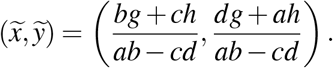

We have the following theorem regarding this equilibrium point.

#### Theorem 6.

*The equilibrium point* 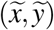 *of the classical Richardson arms race model is either a stable node when ab* > *cd or a saddle point when ab* < *cd*.

*Proof*. The associated Jacobian matrix 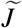 is given by

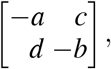

with characteristic equation *λ* ^2^ + (*a* + *b*) *λ* + *ab* − *cd* = 0. We also obtain the following information:

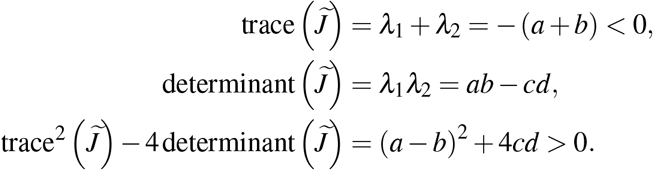

From the final equation, we know that both *λ*_1_ and *λ*_2_ are real numbers. If *λ*_1_*λ*_2_ > 0, which corresponds to *ab* > *cd*, then both eigenvalues are negative, which implies that 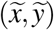 is a stable node. Conversely, if *λ*_1_*λ*_2_ < 0, or *ab* < *cd*, then one of the eigenvalues is positive, whereas the other is negative, e.g., *λ*_2_< 0 < *λ*_1_. This implies that the equilibrium point 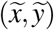 is a saddle point. This completes the proof.

Given a set of initial conditions 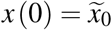 and 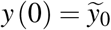, we can also obtain an exact solution to the Richardson arms race initial value problem

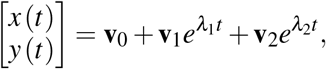

where

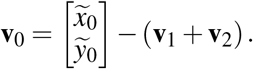

Here, *λ*_1_ and *λ*_2_ are the eigenvalues of the Jacobian matrix 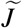

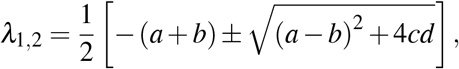

and **v**_1_ and **v**_2_ are the eigenvectors associated with *λ*_1_ and *λ*_2_, respectively.

Figure 11 displays the phase portraits of two examples of the arms race model with different characteristics and the solution curves to their initial value problems. The left panel (Figure 11(a)) corresponds to the system with parameter values *a* = 3, *b* = 4, *c* = 1, *d* = 2, *g* = 3, and *h* = 8, whereas the right panel (Figure 11(b)) admits the parameter values of *a* = 2, *b* = 3, *c* = 3, *d* = 4, *g* = −10, and *h* = −10.

**Figure 11.**
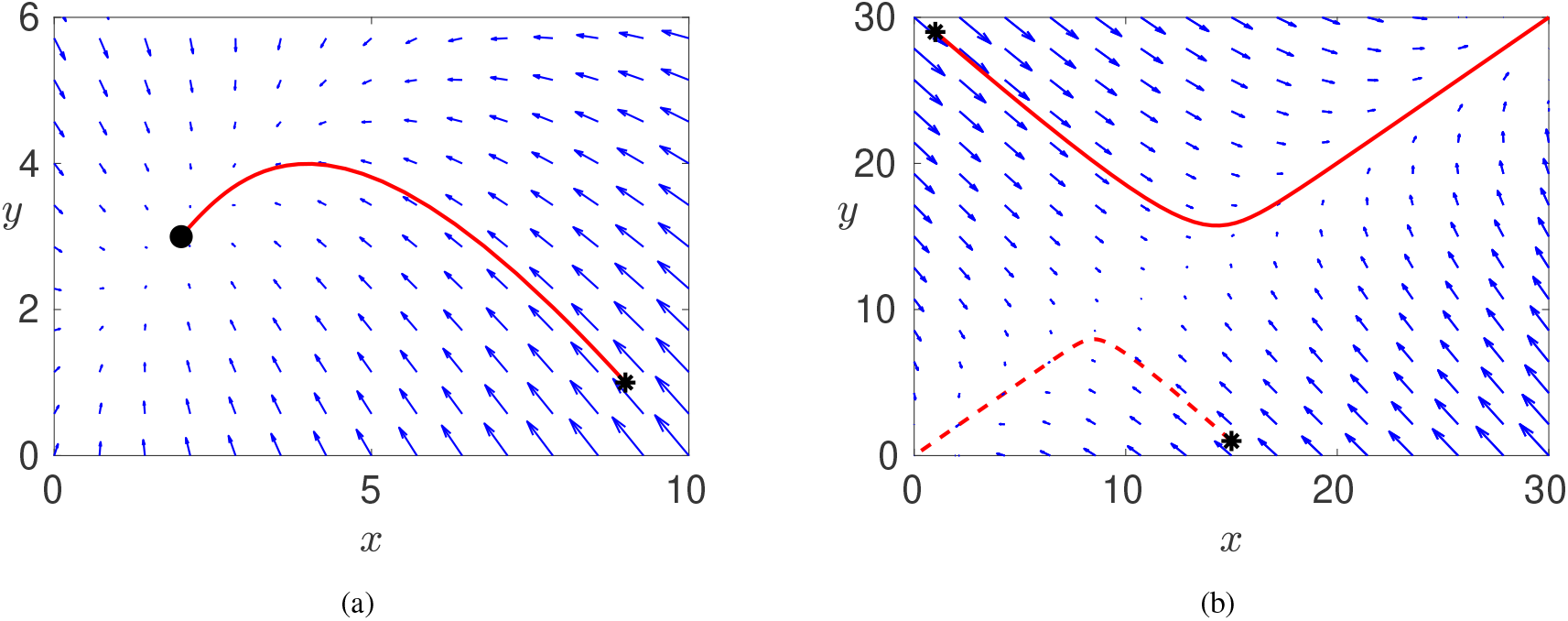
Phase portrait examples of the arms race model together with a solution to the corresponding initial value problem (IVP). (a) An example of a stable equilibrium point where all solutions lead to 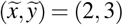 (indicated by a black dot). The red curve denotes the solution to an IVP with initial condition 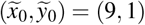 as indicated by a black asterisk. (b) An example where the system has a saddle point at 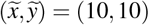. Two distinct initial conditions produce different trajectories and lead to distinct steady state conditions. The dashed red curve has an initial condition at 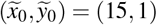, whereas the red solid curve starts at 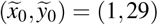. The final points for both situations are not captured in the figure because they are too large in the absolute sense.

As we observe in Figure 11(a), that particular choice of parameters satisfies *ab* = 12 > 2 = *cd*, and thus the equilibrium point 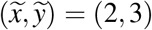 is a stable node. An exact solution to the initial value problem (IVP) with an initial condition 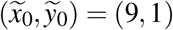 (indicated by a black asterisk) is given by

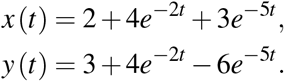

The solution curve is depicted by the solid red curve in Figure 11(a). The steady state solution of the IVP is indeed the critical point, which is indicated by the black dot. Because the equilibrium point is a stable node, the qualitative long-term behavior for such an arms race model will be identical, irrespective of the initial conditions.

Thus, even though the two nations have different proportionality factors for the internal brakes on arms increases, i.e., *a* = 3 vs. *b* = 4, as well as different measures of mutual fear, i.e., *c* = 1 vs. *d* = 2, there is a predictable and fixed point at which the arms levels of both nations will eventually stabilize. This suggests a peaceful and self-correcting dynamic in the arms race, where a balanced state of armament can be achieved and maintained over the long term, without ever-increasing escalation or persistent cycles of buildup and reduction, even though there exist underlying factors of distrust that would persist even if arms expenditures drop to zero, as indicated by the constants *g* = 3 vs. *h* = 8.

Due to a different choice of parameters, which satisfy *ab* = 6 < 12 = *cd*, the phase portrait presented in Figure 11(b) shows different characteristics from the previous case. Instead of a stable node, the equilibrium point 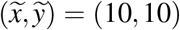 is a saddle node, which is unstable. This means that if the arms levels of the nations deviate even slightly from this precise point, it will move away from it, not back toward it.

A saddle node is characterized by possessing stable and unstable separatrices (or manifolds). On the one hand, if an initial condition starts exactly on the former, then the solution trajectory will approach the equilibrium and converge to the saddle point. On the other hand, the unstable separatrix indicates a specific direction along which solution trajectories would move away from the equilibrium. Even the slightest nudge off the stable manifold will cause the system to be repelled along the unstable manifold.

Figure 11(b) also depicts two solution trajectories emanating from two different initial conditions, as indicated by the black asterisks. Although both trajectories follow the same path along the unstable manifold, both solutions follow different directions and their long-term behaviors end in the opposite direction of the separatrix. The dashed red curve starts with an initial condition of 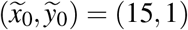 and admits the following solution:

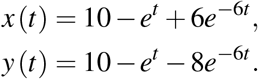

Another solution with a different initial condition of 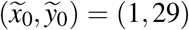 is indicated by the solid red curve, and its analytical solution is given by:

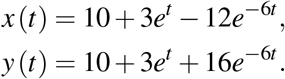

Observe that for the former, the dominant terms in long-term behavior is −*e*^*t*^, whereas in the latter, the dominant term is +3*e*^*t*^ for both *x* and *y*. Thus, we understand why the dashed red curve goes to the third quadrant (both *x* and *y* values are negative) while the solid red curve continues along the separatrix in the first quadrant (both *x* and *y* values are positive). The final points in both cases are not shown in the figure because they are too large in the absolute sense.

From this particular example, we observe that although the equilibrium point at (10, 10) represents a theoretical state where two nations or two universities maintain 10 units of arms or amenities, respectively, this balance is extremely precarious and very sensitive to disturbance. Certainly, their arms or amenities levels would not change if both parties stayed perfectly at this particular point; however, if one party deviates even slightly from (10, 10), e.g., one nation slightly increases its arms to 10.1, or decreases to 9.9, the theoretical model predicts that the dynamics of the system will not bring them back to (10, 10), but will be pushed away instead along the unstable manifold.

This means that such a system is prone to unchecked escalation. Through continuous and accelerating buildup of weapons, such arms race act is unsustainable because it takes an immense drain on resources, increases the risk of catastrophic conflict, and diverts resources from other, more important societal needs. In the amenities arms race in higher education, universities and colleges incessantly increase spending on non-academic amenities. This leads to ever-higher tuition fees, escalating debt for students, and a potential degradation of the core educational mission. Following such a trajectory is certainly unsustainable for students, parents, and the institutions themselves in the long run.

In the context of negative sustainability, the appearance of a saddle point can act as a mathematical warning sign. Unless deliberate and significant interventions are made to alter the fundamental dynamics of the race, such a system may lead to large and unsustainable consequences.

While the classical Richardson arms race model offers valuable insights, real-world interactions often exhibit more complex, non-linear dependencies. To capture such nuances, we can modify the model by allowing the reactions to depend on the arms levels of respective nations, that is, *c* = *c* (*x*) and *d* = *d* (*y*), we can modify the arms race model to become a nonlinear one. For example, by assuming linear functions for *c* and *d*, we obtain a Lotka-Volterra-like model, which is similar to taking functional response of Type I (linear):

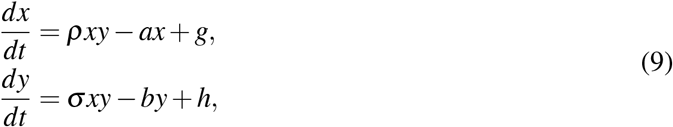

where the coefficients *ρ* and *σ* > 0 are the slope strength for each functional response.

Nevertheless, there exist notable fundamental differences between this modified arms race model and the classical Lotka-Volterra model nonetheless. First, the arms race model includes external factors *g* and *h*, which are usually absent in the Lotka-Volterra model. In the context of predator-prey systems, positive constant terms (*g, h*) would represent *stocking* or *replenishing* the population, while negative constant terms would represent *constant harvesting*. Although it might be relatively rare in the context of ecological systems, this negative harvesting situation can be very beneficial for sustainability.

Second, in the modified Lotka-Volterra model (9), the equation for the prey, that is, for the *x* variable, the terms have the opposite signs compared to the classical one (the standard Lotka-Volterra model). This would create a fundamentally different and quite unusual ecological scenario, but this is a natural and common tendency in the arms race situation. The prey population has a negative intrinsic growth rate and the presence of predators actually supports its growth. In this modified system, the prey population would go extinct without predators, but it can potentially thrive when predators are present at appropriate levels. This represents a form of *obligate mutualism* rather than a predator-prey relationship.

There are other possibilities to modify the arms race model involving nonlinear terms. The body of published literature has many examples of nonlinear arms race models and their applications in social and political sciences (Caspary 1967, Tomochi and Kono 1998, Wallace and Wilson 1978).

## 4 Conclusion

In this chapter, we thoroughly explored the classical Lotka-Volterra predator-prey model, demonstrating its foundational role in understanding complex population dynamics and their profound implications for sustainability, both positive and negative. We analyzed how the model illuminates principles of dynamic equilibrium, stability, and the oscillatory nature of ecological cycles, underscoring the delicate balance essential for sustainable population management. Under ideal conditions, it reveals how predator and prey populations can maintain stable, repeating patterns, thereby avoiding extinction or uncontrolled growth—a key to robust ecosystem stability.

The model further emphasizes the critical role of feedback loops, where predator and prey populations intricately influence each other’s growth and survival. These dynamics offer a microcosm of broader ecological principles, highlighting the interplay of resource availability and natural constraints on species populations. By rigorously understanding these mathematical insights, we can better inform strategies for addressing real-world sustainability challenges, from optimizing natural resource management to fostering effective biodiversity conservation.

Applied within ecological contexts, even simplified models like the Lotka-Volterra system offer profound insights for sustainable predator-prey management. Recognizing and embracing the natural cyclical dynamics of these populations is paramount for designing conservation efforts that foster more resilient ecosystems and ensure long-term resource availability.

Specifically, the ability to predict population peaks and troughs based on these cyclical patterns is invaluable for implementing informed harvesting strategies and establishing appropriate monitoring frequencies. Ultimately, effective and sustainable management hinges on a deep understanding of these complex ecological interactions, empowering wildlife managers to make wise decisions that truly support the health and balance of our planet’s diverse species for generations to come.

Beyond ecological systems, we demonstrated how the Lotka-Volterra framework, particularly through its extension to arms race models, provides crucial insights into competitive dynamics across diverse fields. Whether applied to international conflicts, evolutionary adaptations, or even competition among higher education institutions, these models vividly illuminate how uncontrolled competitive interactions can lead to negative sustainability—outcomes characterized by depletion, self-harm, and the undermining of long-term well-being.

Ultimately, by identifying the critical mechanisms leading to negative sustainability, mathematical modeling offers essential tools for understanding complex systems despite inherent simplifications. Recognizing these dynamics is a vital first step toward addressing the underlying causes of unsustainable behaviors. The profound implication is that by re-directing competitive energy towards genuine and positive collaboration, we can foster a truly sustainable future for all aspects of human and natural systems, transcending the cyclical traps observed in our models.

